# Mechanism for the initiation of co-transcriptional pre-60*S* assembly

**DOI:** 10.64898/2026.05.22.727207

**Authors:** Rafal Piwowarczyk, Sebastian Klinge

**Affiliations:** Laboratory of Protein and Nucleic Acid Chemistry, The Rockefeller University, New York, New York 10065

**Author notes:** Correspondence (S.K.).

## Abstract

Eukaryotic ribosomal large subunit (60*S*) assembly requires an internal transcribed spacer 2 (ITS2) to license both nucleolar and nuclear pre-60*S* assembly intermediates. The underlying molecular mechanisms responsible for nucleation of pre-60*S* assembly, quality control, and installation of ITS2 during co-transcriptional stages remain unknown. Here we report the earliest co-transcriptional assembly intermediates of the eukaryotic 60*S* subunits. Together with biochemical assays, our data reveal the architecture of co-transcriptional pre-60*S* assembly initiation and progression, as well as the molecular logic of an assembly checkpoint. This study highlights an evolutionary solution by which complex RNA folding processes can be parallelized and integrated via biological AND-gating.

## MAIN

In contrast to its bacterial counterpart, eukaryotic ribosome assembly is marked by the presence of both internal and external transcribed spacers, which have been coopted as recruitment platforms for numerous assembly factors^1,2^. During co-transcriptional ribosome assembly, as originally visualized on Miller chromatin spreads, terminal structures are observed, which represent the first ribosome assembly intermediates^3^. However, the mechanism responsible for initiation of co-transcriptional assembly of the large ribosomal subunit remains unknown.

Internal transcribed spacer 2 (ITS2) separates 5.8*S* and 25*S* rRNA, two of the three rRNAs of the eukaryotic ribosomal large subunit (60*S*), and has emerged as key binding site for proteins regulating first nucleolar assembly (via the assembly factor Erb1) and subsequently nuclear assembly (via the assembly factor Nop53)^4–7^.

Biochemical and genetic data show that early co-transcriptional folding events involving pre-60*S* rRNAs (5.8*S*, ITS2, and 25*S*) together with early ribosomal proteins and numerous assembly factors must be controlled in a manner that is distinct from its bacterial counterpart (23*S*), which lacks ITS2. During the last two decades, a set of 12 so-called “A3” factors (including the Brx1-Ebp2, Nop7-Erb1-Ytm1, Rlp7-Nop15-Cic1 modules, ATPases Has1 and Drs1, as well as single proteins Pwp1 and Nop12) have been identified, which mediate early pre-60*S* rRNA processing and co-transcriptional folding events^8–15^. Intriguingly, with the exceptions of Pwp1, Nop12, and Drs1, all of these proteins have been visualized as pre-formed modules in post-transcriptional pre-60*S* particles located close to ITS2, although the mechanisms by which this ensemble of factors nucleates and quality controls co-transcriptional pre-rRNA folding has remained elusive ^5,6,16,17^. A lack of tools to visualize and study these dynamic events has precluded a mechanistic understanding of this essential RNA folding pathway.

Structural studies of bacterial ribosome assembly using both *in vitro* reconstitution and native conditions *in vivo* have revealed a common core of 23*S* rRNA domain I as center for initial folding of the 23*S* rRNA ^18–20^. However, due to evolutionary divergence in rRNA architecture (such as the addition of ITS2) and ensuing emergence of eukaryote-specific ribosomal proteins and assembly factors, these bacterial ribosome assembly intermediates cannot serve as direct blueprints for understanding eukaryotic ribosome assembly (**Extended Data Fig. 1**). This evolutionary divergence necessitates a eukaryote-specific RNA folding pathway that prioritizes early protein-mediated chaperoning and modular control.

Prior studies have revealed that eukaryotic pre-rRNA mimics can be employed to simulate co-transcriptional ribosomal assembly and identify the hierarchy of assembly factor recruitment to pre-rRNA transcripts of defined length^21–24^. Intriguingly, these studies highlight that among the earliest ribosome assembly factors associating with pre-rRNAs of the 60*S*, Pwp1 and Nop12 (PWP1 and RBM34 in humans) have a relatively short residence time, suggesting transient roles during co-transcriptional ribosome assembly. Biochemical studies of these proteins further suggest that they are part of an early assembly factor cluster including other A3 factors^25,26^, that Nop12 binds near the 5.8*S* rRNA^27^, and that both Pwp1 and Nop12 are required for correct folding of 5.8*S* and incorporation of ITS2 ^ref28^.

### Nucleation of co-transcriptional pre-60*S* assembly

To reveal the earliest states of co-transcriptional pre-60*S* biogenesis, we employed tandem affinity purifications to isolate and determine high-resolution structures of particles containing pre-rRNA mimics and specific assembly factors (**Figs. 1,2, Extended Data Figs. 2-4 and Supplementary Figs. 1-8**), which enabled model building across states (**Supplementary Table 1**). Particles containing Pwp1 and pre-rRNA mimics of different lengths (**Supplementary Figs. 1,2,6**) uncovered an equilibrium of the assembly intermediates Pwp1 RNP and Pwp1 RNP*, which were resolved at 2.5 and 2.6 Å respectively (**Fig. 1, Extended Data Figs. 2, 3 and Supplementary Figs. 6,7**). We further identified a minimal pre-rRNA mimic, which is necessary and sufficient for the formation of Pwp1 RNPs. Strikingly, the formation of the Pwp1 RNP is independent of the presence of ITS1, 5.8S rRNA, and ITS2 and represents the smallest ribosome assembly intermediate determined to date (**Extended Data Fig. 4, Supplementary Fig. 2**). The Pwp1 RNP further highlights the shift towards a protein-orchestrated ribosome assembly pathway in eukaryotes where pre-rRNA is chaperoned very early on so that the resulting RNP is significantly smaller than even the earliest visualized bacterial ribosome assembly intermediates, which only contain a few ribosomal proteins (**Extended Data Fig. 5**) ^18,23^. With only 136 nucleotides of domain I of the 25*S* rRNA acting as initial binding platform, the Pwp1 RNP can be divided into two halves with one half containing three early ribosomal proteins (Rpl8, Rpl15, and Rpl36) and the second half containing ribosome assembly factors (Brx1-Ebp2, and Pwp1) (**Fig. 1a, b**). While the three ribosomal proteins and the Brx1-Ebp2 constitutive dimer act as rigid chaperones of pre-ribosomal RNA, Pwp1 acts as a central nexus that connects all components within the RNP. Key interactions by Pwp1 are largely mediated via peptides that can undergo conformational changes to alternate between Pwp1 RNP and Pwp1 RNP* (**Extended Data Figs. 2,3**). Many of the observed interactions within Pwp1 RNPs are consistent with AlphaFold predictions, which additionally suggest an interaction involving a large peptide of Nop12 with Pwp1 (**Extended Data Fig. 6**) ^29^.

**Fig. 1:**
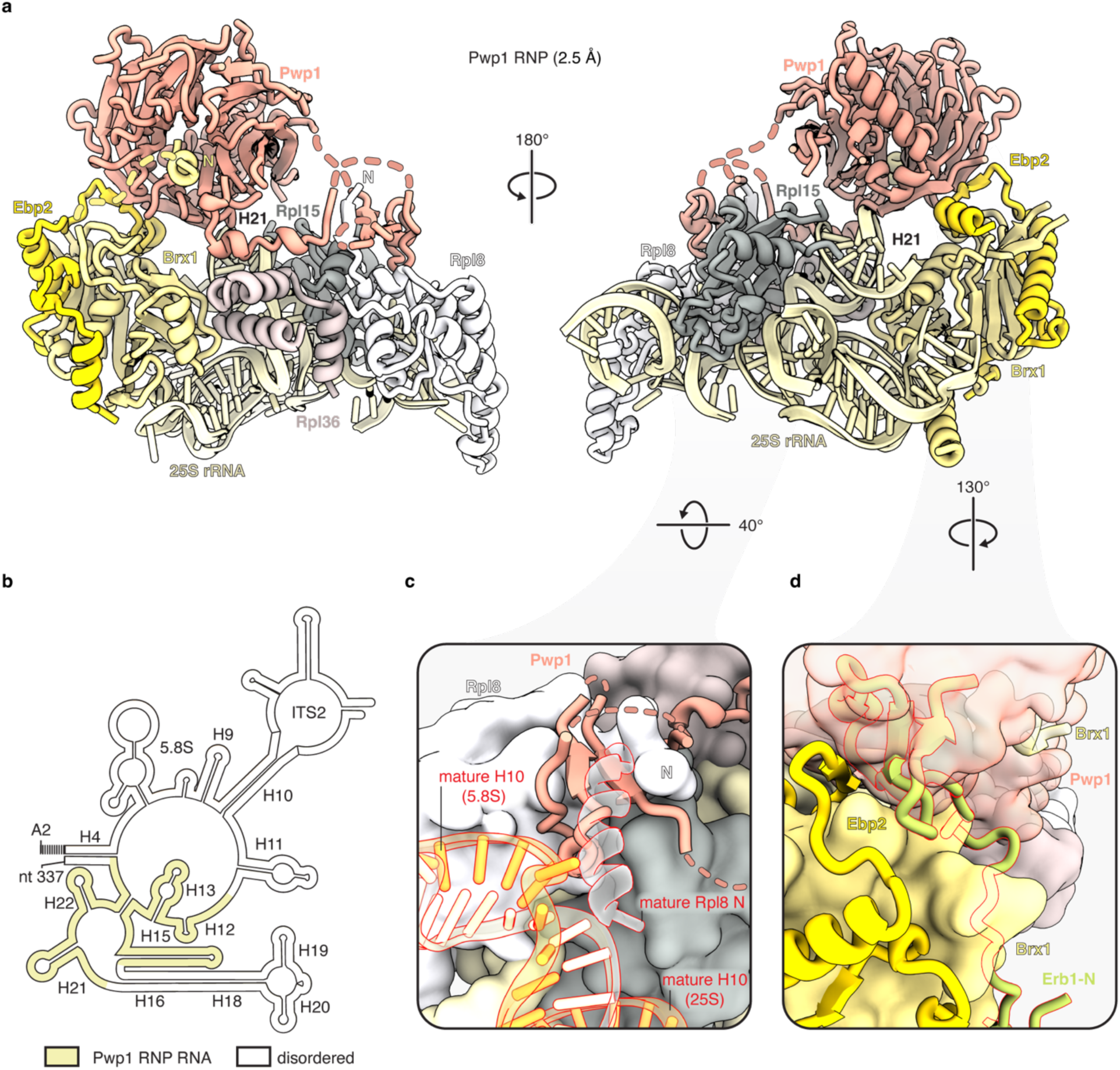
Nucleation of co-transcriptional pre-60*S* assembly. (a) Cryo-EM reconstruction of the Pwp1 RNP at 2.5 Å resolution shown in two orientations. The RNP consists of a 136-nucleotide segment of 25S rRNA domain I chaperoned by three early ribosomal proteins (Rpl8, Rpl15, Rpl36) and an assembly factor module comprising the Brx1-Ebp2 constitutive dimer and Pwp1. Pwp1 serves as a central nexus, connecting the protein and RNA components, nucleating the assembly. (b) Secondary structure diagram of the A2®337 rRNA mimic. Regions ordered within the Pwp1 RNP are highlighted in yellow. (c) Detailed view of the Pwp1-mediated steric checkpoint at the Heli× 10 (H10) docking site. The interaction with Pwp1 prevents the Rpl8 N-terminal tail from reaching its mature conformation and sterically obstructs the formation of the H10 RNA duplex between the 5.8S and 25S rRNAs, effectively stalling maturation at this co-transcriptional stage. (d) Structural basis for the mutual exclusivity between Pwp1 and the nucleolar licensing factor Erb1. Pwp1 occupies a binding site on the surface of the Brx1-Ebp2 dimer that overlaps with the binding site for the Erb1 N-terminal (Erb1-N; green with red outline).

**Fig. 2:**
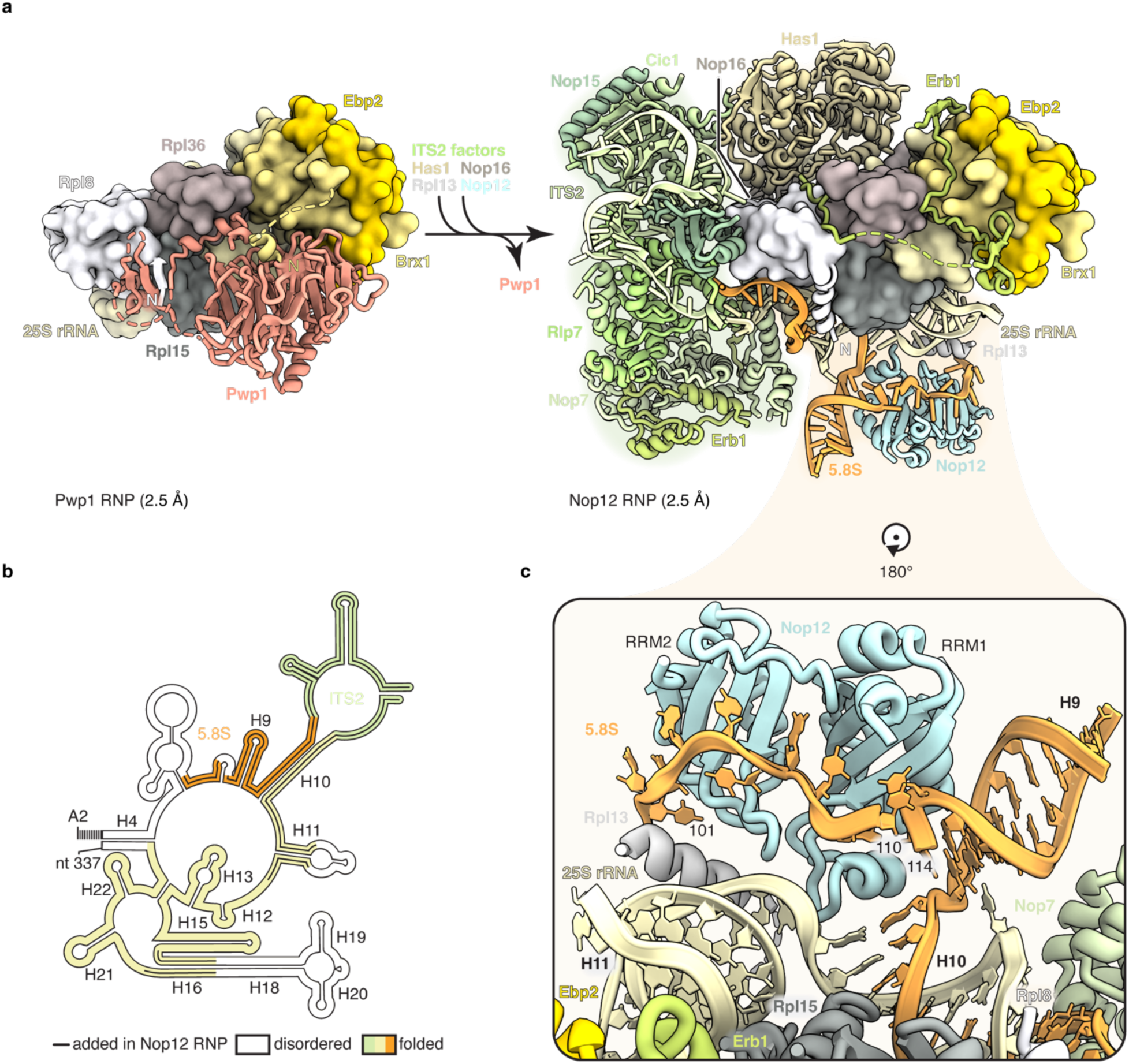
Structural basis for 5.8*S* and ITS2 incorporation. (a) Structural transition from the Pwp1 RNP to the Nop12 RNP (both at 2.5 Å resolution). This maturation step is marked by the departure of Pwp1 (coral), the incorporation of ITS2 with associated A3 factors Has1, Cic1, Nop15, Rlp7, Nop7, and Erb1 (shades of green/tan), ordering of helix H16 stabilized by Nop16, alongside the installation of 5.8*S* rRNA by Nop12 (light blue) and ribosomal protein Rpl13 (light grey). (b) Secondary structure diagram of the rRNA mimic highlighting structural progression. rRNA elements newly ordered in Nop12 RNP, including the 5.8*S* rRNA (orange) and the ITS2 (green), are indicated. Helices H9-11, which flank the ITS2, are now fully formed. (c) Detailed view of the Nop12 interface with the 5.8*S* rRNA. Nop12 utilizes the two RNA recognition motifs domains (RRM1 and RRM2) to provide sequence-specific binding of a single-stranded region of the 5.8*S* rRNA. Together with Rpl13, Nop12 stabilizes the pre-mature folding intermediate of the 5.8*S* rRNA.

The structure of the Pwp1 RNP highlights two regulatory roles of Pwp1 during co-transcriptional stages of pre-60*S* biogenesis. First, the ability of Pwp1 to bring together five additional proteins ensures rapid stabilization of a small RNA nucleation point. Second, Pwp1 binding precludes premature formation of interactions observed in subsequent states. Here both the incorporation of ITS2 via heli× 10 (H10), a key RNA duplex between the 5.8*S* and 25*S* rRNAs near Rpl8, and the binding of the nucleolar licensing factor Erb1 near the Brx1-Ebp2 dimer are prevented by Pwp1 (**Fig. 1c,d and Extended Data Figs. 2,3**). This mutual exclusivity between Pwp1 and productive pre-rRNA compaction of domain I and ITS2 rationalizes the transient nature of Pwp1 association and highlights its stalling function during early co-transcriptional pre-60*S* biogenesis.

### Incorporation of ITS2 and 5.8*S* into co-transcriptional pre-60*S* intermediates

Our data show that RNA chaperoning is a modular and autonomous process. Here pre-rRNA truncation experiments highlight that the elimination of the 5.8*S* rRNA results in the absence of Nop12, suggesting that Nop12 directly chaperones parts of the 5.8*S* rRNA (**Extended Data Fig. 4**).

By using Nop12 and a pre-rRNA mimic as affinity baits, we obtained the 2.5 Å resolution structure of the Nop12 RNP, which is dramatically different from the Pwp1 RNP (**Fig. 2a, Extended Data Fig. 3 and Supplementary Figs. 3,4**). The incorporation of ITS2 changes the overall architecture of this particle and is associated with the partial ordering of the 5.8S and parts of domain I of the 25S rRNA (helices 9-11) (**Fig. 2b,c**). The integrated ITS2 establishes a binding site for the nucleolar licensing factor Erb1, which in turn provides binding sites for the DEAD-box ATPase Has1 as well as the assembly factor Nop16, stabilizing helix 16 (H16) in a bent conformation. Importantly, bending of H16 is strictly required for subsequent states of assembly where extensions of this helix mediate key contacts between domains I, II, and fully folded 5.8*S* rRNA (**Extended Data Fig. 7**). Within the Nop12 RNP, sequence specific recognition of a single-stranded region of the 5.8*S* rRNA is provided by Nop12. Nop12 and ribosomal protein Rpl13 stabilize a pre-mature folding intermediate of the 5.8*S*, which is incompatible with the Rpl35-bound mature 5.8S visualized in later assembly states (**Fig. 2b and Extended Data Figs. 3l,7b,c**). The transition from the Pwp1 RNP towards the Nop12 RNP rationalizes the early functions and essential roles of the A3 factors during early pre-60S biogenesis^30^. These data further reveal that the insertion of ITS2 requires separate chaperoning of the 5.8S and stepwise installation of H16, which is not necessary in bacteria (**Extended Data Fig. 7**).

### Identification of a ribosome assembly checkpoint

The compositional and conformational changes observed during the transition from the Pwp1 RNP to the Nop12 RNP suggest that several inputs are required to trigger unidirectional maturation. RNA conformational changes that could have an impact on this transition include the incorporation of ITS2, the binding site for several assembly factors, as well as H16, which is stabilized in a bent conformation underneath Has1. Protein elements that could affect this transition are the ATPase Has1, as well as the N-terminal segment of Erb1 (Erb1-N), which is mutually exclusive with Pwp1 (**Fig. 1c, 3a-c and Extended Data Fig. 3e,j,o**). Purifications of RNPs containing pre-rRNA mimics lacking either H16 or ITS2 showed that their elimination results in the dramatic reduction of ITS2-associated factors Erb1 and Ytm1 (**Fig. 3d**). To verify the structural consequences of ITS2 and H16 removal, purified mutant and full-length samples were further analyzed using cryo-EM and processed together. While the wild-type pre-rRNA mimic could form both Pwp1 and Nop12 RNPs, mutants lacking either H16 or ITS2 only gave rise to Pwp1 RNPs and were unable to transition to a Nop12-bound state (**Supplementary Fig. 6**). These data suggest that the ability to correctly incorporate ITS2 and stabilize a bent H16 are integral parts of an early ribosome assembly checkpoint. As both events are further supported by the installation of proteins (Erb1 for ITS2 and Has1 for H16, respectively), their contributions in Nop12 RNP formation were investigated.

**Fig. 3:**
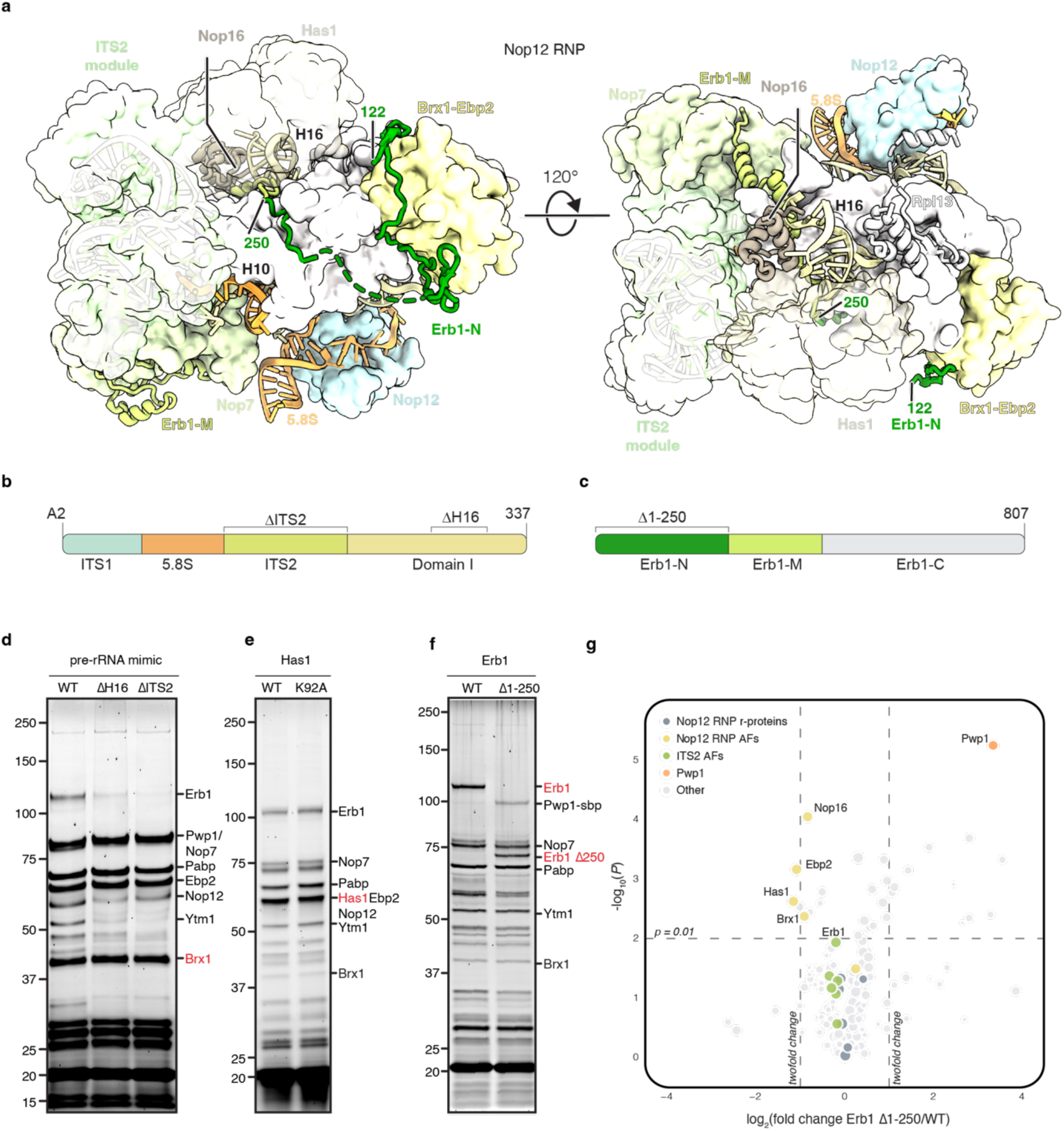
Molecular logic of a co-transcriptional assembly checkpoint. (a) Two orientations of the Nop12 RNP structure (2.5 Å) highlighting the trajectory of Erb1 and its N-terminal (Erb1-N, dark green) and middle (Erb1-M, light green) segments. Erb1 wraps around the intermediate, interacting with the Brx1-Ebp2 constitutive dimer, stabilizing Helix 16 (H16) alongside Has1 and Nop16, and contacting Nop7 within the ITS2. The ITS2 module, Nop12, and 5.8S rRNA are colored as in Fig. 2. (b) Schematic of the A2®337 pre-rRNA mimic used for biochemical and structural analyses. Brackets indicate the specific regions targeted for deletion (ΔITS2 and ΔH16). (c) Schematic of Erb1, highlighting the N-terminal segment (Erb1-N, residues 1-250, dark green), the middle domain (Erb1-M; light green), and the C-terminal domain (Erb1-C; grey). (d-f) SDS-PAGE analysis of affinity-purified RNPs. Complexes were isolated via a two-step tandem purification utilizing a protein assembly factor as the primary bait (red text, Brx1 (d), Has1 (e), and Erb1 (f)) followed by the rRNA mimic enrichment. (d) Analysis of particles obtained using pre-rRNA mimic mutants. Absence of either H16 or ITS2 results in a significant reduction of ITS2-associated factors Erb1 and Ytm1. (e) Compositional analysis of particles containing wild-type Has1 or the catalytically inactive Has1(K92A) mutant showing no difference in assembly factor composition between conditions. (f) Analysis of pre-ribosomal particles obtained in Erb1 truncation experiment. Deletion of the Erb1(Δ1-250) N-terminal segment leads to a marked increase in Pwp1 levels while other factors remain similar. (g) Comparative mass spectrometry analysis of RNPs purified with Erb1(Δ1-250) mutant or WT. Pwp1 (orange) is enriched 10-fold in the mutant, while other assembly factors remain at a similar level. Three biological replicates were analyzed in each condition. *P* values were determined by a two-tailed Student’s *t*-test, and a *P* value of 0.01 was chosen as the cut-off for statistical significance.

A catalytically inactive Has1(K92A) mutant can be co-enriched with Erb1-containing particles, highlighting that the ATPase activity of Has1 is not required for ITS2 incorporation and Erb1 association during co-transcriptional pre-60*S* assembly (**Fig. 3e**). These findings are consistent with prior biochemical data indicating that the ATPase activity of Has1 is not required for its initial association^31^. Separately, the structure of the Nop12 RNP suggests that the installation of the Erb1 N-terminal segment (Erb1-N) is incompatible with the presence of Pwp1 and Erb1-N likely acts as gatekeeper preventing Pwp1 reassociation following successful ITS2 incorporation. As hypothesized, the deletion of the Erb1 N-terminal segment (Δ1-250) resulted in a 10-fold increase of Pwp1 in these particles, solidifying the role of the Erb1 N-terminal segment in clearing of Pwp1 to pass the checkpoint. The N-terminal Erb1 truncation also results in a slight reduction in Has1, Nop16 and Brx1-Ebp2 as these proteins are not stabilized as much (**Figs. 3a,f,g**). These data suggest that upon incorporation of ITS2 and bending of H16, the N-terminal segment of Erb1 directly competes with Pwp1 for a binding site on the surface of the Brx1-Ebp2 complex and is responsible for the completion of the co-transcriptional ribosome assembly checkpoint.

### An emerging model for the initiation of co-transcriptional pre-60*S* assembly

Our data now allow us to propose a model that rationalizes two decades of biochemical and genetic observations. During co-transcriptional RNA folding, three elements are chaperoned in a modular and parallelized way. These include parts of domain I that are stabilized within the Pwp1 RNP by ribosomal proteins (Rpl8, Rpl15, and Rpl36) and assembly factors (Brx1-Ebp2 and Pwp1), parts of the 5.8*S* that are chaperoned by Nop12, and ITS2 that is stabilized by dedicated assembly factors (Cic1, Nop15, and Rlp7). The intramolecular connectivity of these early modules is ensured both by the underlying RNA, its ability to bring segments in proximity via base-pairing between 5.8*S* and 25*S* elements, and the ability of protein elements to weakly interact (**Fig. 4a**). The molecular underpinnings of the assembly checkpoint involve an initial installation of ITS2, forming the basis for subsequent events, which can be thought of as mutually dependent and reinforcing equilibria (**Fig. 4b**). Here the stabilization of ITS2 by the Erb1-Ytm1-Nop7 complex^9^ results in the formation of an Erb1-stabilized platform onto which Has1 and Nop16 can assemble to chaperone H16 in a bent conformation. The completion of this step provides the N-terminal segment of Erb1 with the opportunity to outcompete and thus release Pwp1 from this intermediate to complete the checkpoint. Docking of the 5.8*S* via Nop12 is further stabilized by the association of Rpl13, thereby completing the transition towards the Nop12 RNP (**Fig. 4c**). Subsequent formation of domain II then results in a Noc1-Noc2 RNP, which was described previously^17^. The high-resolution structure of the Nop12 RNP now also rationalizes prior low-resolution data for domain I of the Noc1-Noc2 RNP, suggesting that Nop12 can co-exist with Noc1-Noc2 containing particles in which domains I and II have not yet been joined^17^. These early transitions provide an unprecedented view of how eukaryotic co-transcriptional pre-60S assembly is set up to ensure stepwise and unidirectional progressions that accompany the specific readout of key interactions between the 5.8S and 25S rRNA via helices H2,H4, and H10 (**Fig. 4d**).

**Fig. 4:**
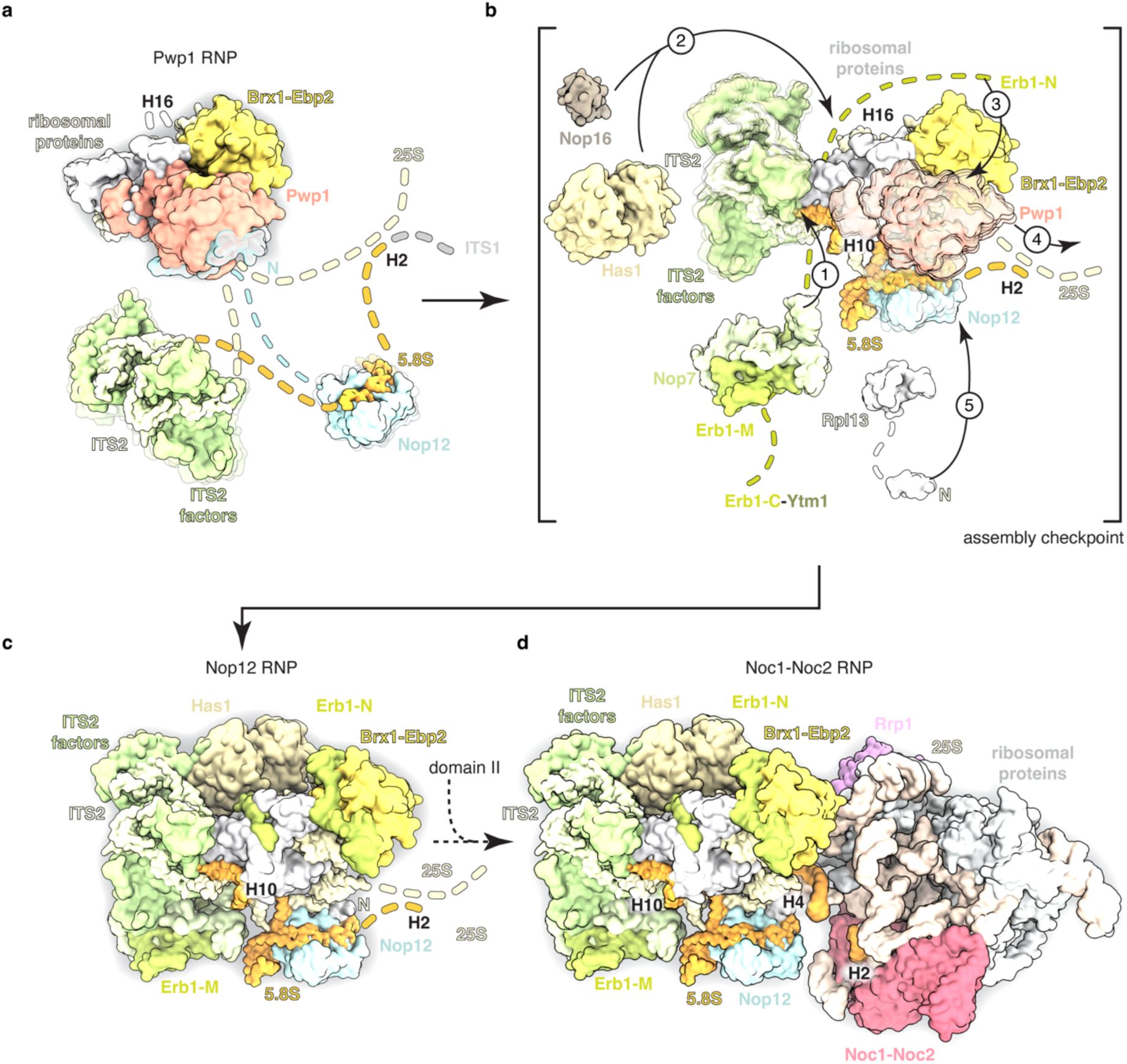
Mechanism of co-transcriptional pre-60S assembly. (a) Modular and parallelized co-transcriptional assembly of early pre-60S. During co-transcriptional biogenesis, three distinct RNA-protein modules are chaperoned independently: (i) domain I of the 25S rRNA within the Pwp1 RNP, (ii) the 5.8S rRNA by Nop12, and (iii) the ITS2 by early binders Cic1, Nop15, and Rlp7. Connectivity between these modular units is facilitated by nascent base-pairing and weak protein-protein contacts (dashed lines). (b) Molecular logic of the biological AND-gate checkpoint. The assembly checkpoint integrates multiple structural inputs to ensure unidirectional maturation: (1) formation of the H10 RNA duplex between 5.8S and 25S rRNA and recruitment of the Erb1-Ytm1-Nop7 complex leading to incorporation of completed ITS2 module. (2) stabilization of H16 into a bent conformation by Has1 and Nop16 and (3,4) the displacement of Pwp1 by the Erb1 N-terminal segment (Erb1-N). Finally, (5) docking of Nop12-5.8S rRNA through an interface created by Rpl13. (c) Successful clearance of the checkpoint results in the formation of the Nop12 RNP, where the 5.8S and 25S rRNAs are brought in close proximity by H10 and Nop12, increasing the probability of H4 and H2 base-pairing necessary for the formation of the next intermediate (d) The subsequent folding of domain II leads to the formation of the Noc1-Noc2 RNP.

### Conclusions

The evolution of pre-rRNA spacers (5’ETS and ITS2) as scaffolding systems for eukaryotic ribosome assembly factors has dramatically changed the nature and complexity of the underlying RNA folding pathways. Where in bacteria the 23*S* rRNA can form an early core intermediate of domain I, which includes several ribosomal proteins, the presence of ITS2 in eukaryotes imposes both RNA folding hurdles and necessitates mechanisms to ensure independent folding of the 5.8*S*, ITS2 and parts of domain I, which are integrated by the A3 factors via biological AND-gating. Here the use of multiple inputs enables both rapid parallelized RNA folding of individual units without compromising overall quality control since an early ribosome assembly checkpoint identified in this study only allows correctly folded RNPs to be licensed for subsequent Erb1-mediated nucleolar assembly. The A3 factors are thus an ensemble of early ribosome assembly factors that compensate the emergence of ITS2 and a separate 5.8*S* rRNA by orchestrating this early ribosome assembly checkpoint. The principles highlighted by this study are broadly applicable to large RNA-protein complexes in which independent folding events coupled with checkpoints provide evolutionary solutions to complex RNA folding processes.

## METHODS

### Yeast strains

Strains were derived from *Saccharomyces cerevisiae* BY4741 expressing an MCP-GFP fusion protein under the control of a galactose-inducible *GAL10* promoter^21^. Assembly factors (Pwp1, Nop12, and Brx1) were endogenously tagged at the C-terminus with a cleavable mCherry tag via PCR-based homologous recombination. For Western blot analysis, specific factors were further tagged with a C-terminal streptavidin-binding peptide (sbp). Genomic integrations were validated by PCR and Sanger sequencing.

Yeast were transformed with pESC-URA plasmids encoding pre-60S rRNA mimics under the *GAL1* promoter. In experiments assessing the impact of factor mutations, strains were co-transformed with a pESC URA based rRNA mimic and an additional centromeric pRSII415-LEU plasmid carrying N-terminally tagged Erb1 (NLS-mCherry-TEV-linker) or C-terminally tagged Has1 (linker-TEV-mCherry) under their native promoters.

### Purification of pre-60*S* intermediates

Cultures were grown in synthetic drop-out (SD) medium containing 2% galactose at 30 °C for 16-18 hours or until reaching an optical density at 600 nm (OD_600_) of 2. Cells were harvested (3,000 x g, 10 minutes, 4 °C), washed twice with ice-cold ddH_2_O, and once with ddH_2_O containing protease inhibitors (E-64, pepstatin, PMSF) before flash-freezing in liquid nitrogen.

Cells were lysed via four cycles of cryogenic grinding (Retsch PM100) and resuspended in Buffer A (50 mM HEPES pH 7.5, 150 mM KCl, 5 mM MgCl_2_, 1 mM DTT, 0.02 % NP-40, and 0.2 % *n*-Octyl β-D-thioglucopyranoside) with protease inhibitors (PMSF, pepstatin, E-64). The lysate was centrifuged (4 °C, 40,000 x g, 30 min), and the supernatant was incubated with anti-mCherry nanobody-coupled NHS-Sepharose beads for 1 hour at 4 °C. The beads were washed in buffer A and bound complexes were eluted via TEV cleavage for 1 hour at 4 °C. Samples selected for cryo-EM analysis were biotinylated (0.5 mM Sulfo-NHS-Biotin, Thermo Fisher Scientific) for 30 min at room temperature and quenched with 10 mM Tris-HCl (final concentration). The sample was then incubated with anti-GFP nanobody beads for 1 hour at 4 °C. The beads were washed three times in Buffer A and three times in Buffer B (50 mM HEPES pH 7.5, 150 mM KCl, 5 mM MgCl_2_, 1 mM DTT, and 0.2 % *n*-Octyl β-D-thioglucopyranoside) before elution via 3C protease cleavage for 45 min on ice.

### Cryo-EM grid preparation and data acquisition

Quantifoil Au R3.5/1 grids with 2-nm ultrathin carbon (EMS) were glow-discharged for 30 s and transferred to a Vitrobot Mark IV (FEI Company) for vitrification (100% humidity, 10 °C). Sample (3.5 µL) was applied to the grid and following a 3 min wait time, the grid was blotted (blot force 6, blot time of 6 s) and plunged into liquid ethane.

Data were collected on a Titan Krios (FEI) operating at 300 kV with an energy filter (20 eV slit width) and a K3 detector (Gatan). Movies were acquired across a defocus range of −0.7 to −1.7 µm at a super-resolution pixel size of 0.54 Å using SerialEM^32^. 50 frames were collected during a 2 s exposure time with a dose of ∼30 e^−^/px/s resulting in a total dose of ∼50.5 e^−^/Å^2^. A multi-shot protocol was used following a 2×2 pattern at each stage position with 11 micrographs per hole.

### Cryo-EM data processing

All datasets were processed following the same workflow in cryoSPARC (v.4.6.0)^33^. Micrographs were 2x binned (1.08Å final pixel size), motion-corrected, and subjected to patch-CTF estimation. To account for beam tilt and higher-order aberrations, micrographs were separated into unique optics groups based on their multi-shot hole position. Particles were picked using a blob picker and extracted with a 300 px box size. Following an extensive 2D-classification round, a set of classes with distinct views was used for *ab initio* map generation and to train a particle picking model using crYOLO (v.1.9.9)^34^. Particles picked by both methods were combined and de-duplicated. The resulting particle stack underwent 5-7 rounds of heterogeneous refinement using 20 Å low-passed *ab initio* maps as references. High-quality classes from the final round of the heterogenous refinement were combined and refined via Non-Uniform (NU) refinement^35^ with CTF-refinement^36^ enabled. The particle stack and volume were then polished via Reference Based Motion Correction, followed by re-refinement to generate a final consensus map. This map was subsequently 3D-classified to resolve any conformational heterogeneity, and the best classes were refined using NU-refinement.

### Generation of focused and composite maps for model building

Local refinement was guided by observed flexibility. Masks for local refinement were generated based on published or AlphaFold3 predicted models^29^ using the molmap command (40 Å resolution) in ChimeraX^37^ and subsequently dilated in cryoSPARC. Each segment underwent a single round of local refinement with a narrow Gaussian prior and recentering at each iteration.

All maps were rescaled to a calibrated pixel size of 1.06 Å/px and sharpened via phenix.auto_sharpen^38^, with the resolution parameter set to values at the GSFSC = 0.143 threshold. Focused maps were combined using phenix.combine_focused_maps to generate a final composite map. Local resolution for all consensus and focused maps was estimated within cryoSPARC.

### Model building and refinement

Atomic models were constructed using a combination of existing cryo-EM structures^17^ (PDB 8E5T), AlphaFold3 predictions, and *de novo* building. Initial models were manually adjusted in ISOLDE^39^ and Coot^40^ to fit the density. The final models were refined against the composite maps using phenix.real_space_refine (v.1.19), with secondary structure restraints applied to both protein and RNA chains. Model validation and refinement statistics are summarized in Table 1. All structural analyses and illustrations were generated in UCSF ChimeraX v1.10.1.

### Comparative mass spectrometry

Purified RNP samples were reduced and alkylated before overnight acetone precipitation. Pellets were recovered by centrifugation, dried, and resuspended for subsequent trypsin and Lys-C digestions. Digestion was quenched with neat trifluoroacetic acid (TFA), and the resulting peptides were purified via solid-phase extraction. Samples were analyzed by liquid chromatography-tandem mass spectrometry (LC-MS/MS) using an Orbitrap Fusion Lumos Tribrid mass spectrometer operated in high-resolution/high-mass-accuracy mode. Peptides were separated over a 70-minute analytical gradient on a 12-cm pulled emitter column (C18 reversed-phase). Raw data were processed using MaxQuant^41^ (v2.6.6.0) to generate Label-Free Quantification (LFQ) and iBAQ values. Searches were performed against the *Saccharomyces cerevisiae* UniProt database concatenated with the MCP sequence and common contaminants. MaxQuant was used to produce LFQ and iBAQ values. Data were normalized using the MaxLFQ algorithm and log2-transformed. Proteins identified as contaminants were removed, and filtering was applied to retain only those proteins identified in at least two out of three replicates within at least one experimental group. Missing values were imputed to facilitate statistical analysis via a two-tailed Student’s t-test.

## Data availability

Original uncropped images of all SDS-PAGE gels used in this study are provided as **Supplementary Fig. 8**. The cryo-EM density maps and atomic coordinates have been deposited in the Electron Microscopy Data Bank (EMDB) and the Protein Data Bank (PDB). **Pwp1 RNP**: Overall map (EMD-75519), Pwp1 focused map (EMD-75520), Composite map (EMD-75577), and Atomic model (PDB 10ZY). **Pwp1 RNP**\*: Overall map (EMD-75539), Pwp1 focused map (EMD-75540), Rpl8 focused map (EMD-75541), Composite map (EMD-75578), and Atomic model (PDB 10ZZ). **pre-Nop12 RNP**: Overall map (EMD-75542), Focused maps for Cic1 (EMD-75543), Brx1 (EMD-75544), Has1 (EMD-75545), and Nop7 (EMD-75546), and Composite map (EMD-75579). **Nop12 RNP**: Overall map (EMD-75547), Focused maps for Cic1 (EMD-75548), Brx1 (EMD-75549), Has1 (EMD-75550), Nop7 (EMD-75552), and Nop12 (EMD-75553), Composite map (EMD-75580), and Atomic model (PDB 11AA). Raw unaligned multi-frame movies and aligned micrographs have been deposited in the Electron Microscopy Public Image Archive: EMPIAR-XXXX. Source data for the comparative mass spectrometry are provided with the paper as a Source Data file. Materials are available from S. K. upon reasonable request under a material transfer agreement with the Rockefeller University.

## Acknowledgements

We would like to thank M. Jenkyn-Bedford for his contributions to sample preparation and cryo-EM data processing, and M. Ebrahim, J. Sotiris and H. Ng at the Evelyn Gruss Lipper Cryo-Electron Microscopy Resource Center at The Rockefeller University (RRID:SCR_021146) for assistance with grid screening and data collection. Mass spectrometry data were generated by the Proteomics Resource Center at The Rockefeller University (RRID:SCR_017797) using instrumentation funded by the Sohn Conferences Foundation and the Loena M. and Harry B. Helmsley Charitable Trust, and we particularly thank H. Molina, C. Peralta and S. Heissel for assistance. We also thank members of the Klinge laboratory for critical reading of this manuscript.

## Author information

Laboratory of Protein and Nucleic Acid Chemistry, The Rockefeller University, New York, NY, USA

Rafal Piwowarczyk, Sebastian Klinge

## Contributions

S.K. and R.P. conceived the study, designed the experiments and analyzed data. R.P. performed all experiments, including strain generation, complex purification, cryo-EM data processing and model building. All authors wrote and edited the manuscript.

## Ethics declarations

The authors declare no competing interests.

## Extended Data

**Extended Data Fig 1.**
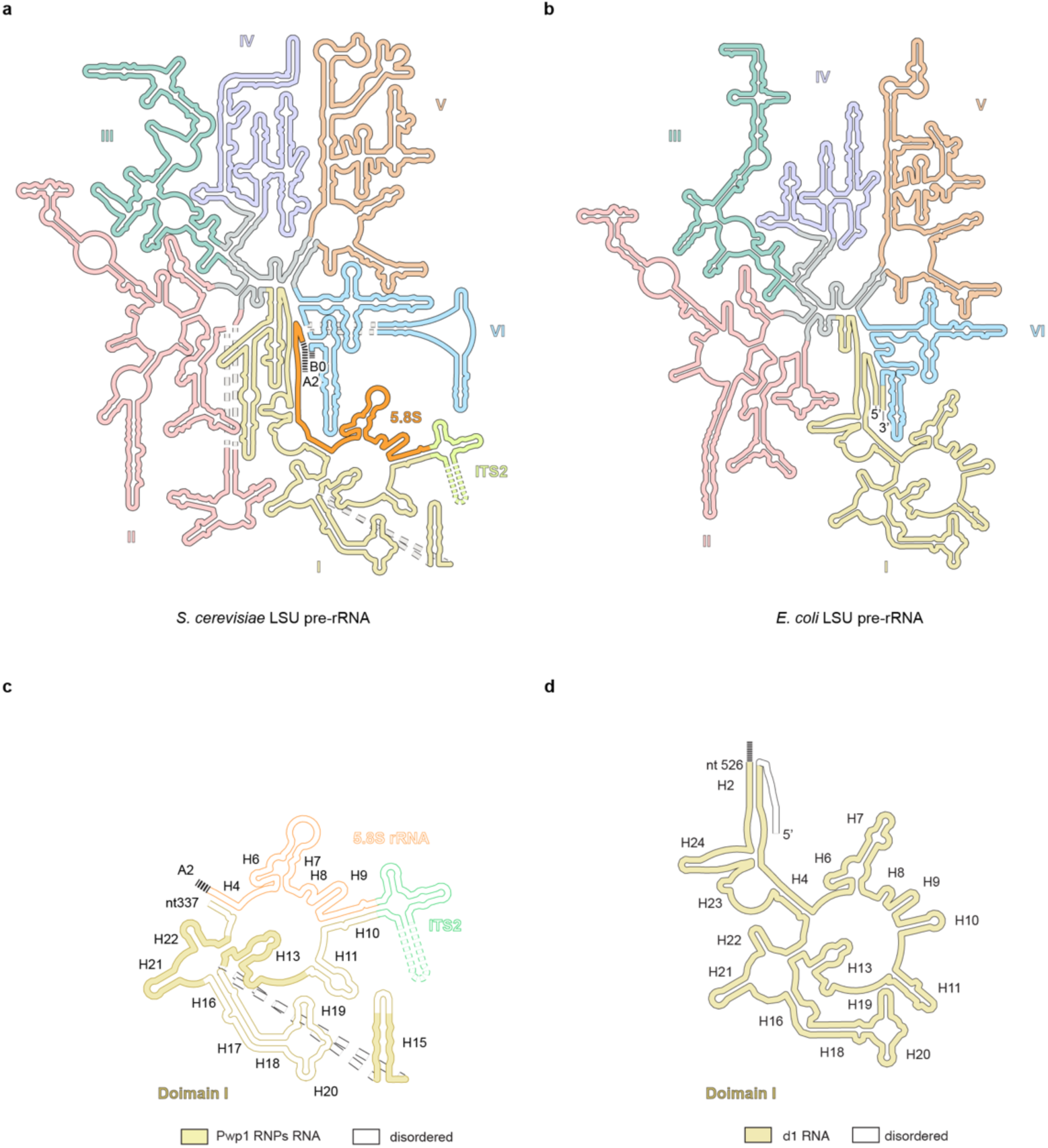
Evolutionary divergence of large ribosomal subunit architecture and the eukaryotic-specific nucleation core. (a, b) Comparison of secondary structures of the LSU pre-rRNA from *S. cerevisiae* (a) and *E. coli* (b). rRNA domains (I-VI) are color-coded to illustrate conserved topological organization. The eukaryotic precursor is distinguished by further rRNA expansion and the insertion of internal spacer ITS2 (green). Notably, the incorporation of ITS2 sequence resulted in significant divergence of LSU rRNA architecture between prokaryotic and eukaryotic species, separating the 5.8S (orange) and 25S rRNAs. Secondary structure diagrams adapted from Petrov et al.^42^ (c, d) Comparison of ordered rRNA scope in the earliest described LSU assembly intermediates between species. (c) The eukaryotic Pwp1 RNP (this study) represents the smallest co-transcriptional nucleation core, where only 136-nucleotide segment of Domain I (yellow) is ordered. (d) The prokaryotic d1 intermediate (PDB: 8C9C) utilizes the entire Domain I for initial 23*S* rRNA folding.

**Extended Data Fig 2.**
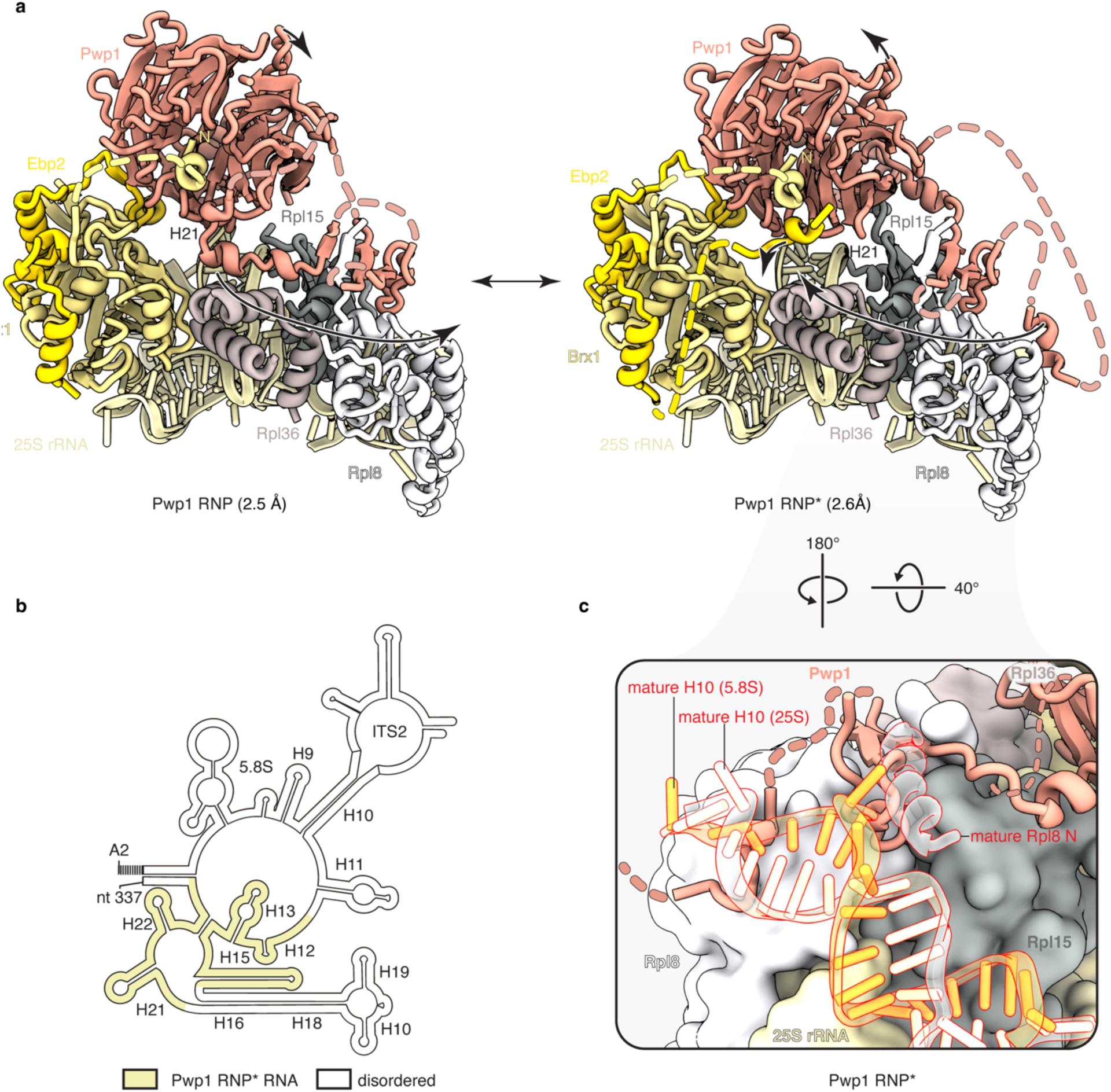
Conformational equilibrium of Pwp1 RNPs. (a) Comparison of the atomic models for Pwp1 RNP (2.5 Å) and Pwp1 RNP* (2.6 Å). The two intermediates exist in a conformational equilibrium defined by a concerted rotation and displacement of the Pwp1 β-propeller domain and its internal loop (black arrows) relative to the rRNA core. (b) Secondary structure representation of the A2→337 rRNA mimic in the Pwp1-RNP* state. Ordered rRNA segments are indicated in yellow. (c) Detailed view of the Pwp1-Rpl8 interface in the Pwp1 RNP* (related to Fig. 1c) illustrating the mechanism of the Pwp1-mediated steric checkpoint. The presence Pwp1 prevents the Rpl8 N-terminal tail from reaching its mature conformation (red) and sterically obstruct the docking of the H10 composed of both 5.8S and 25S rRNA (yellow and white respectively).

**Extended Data Fig. 3.**
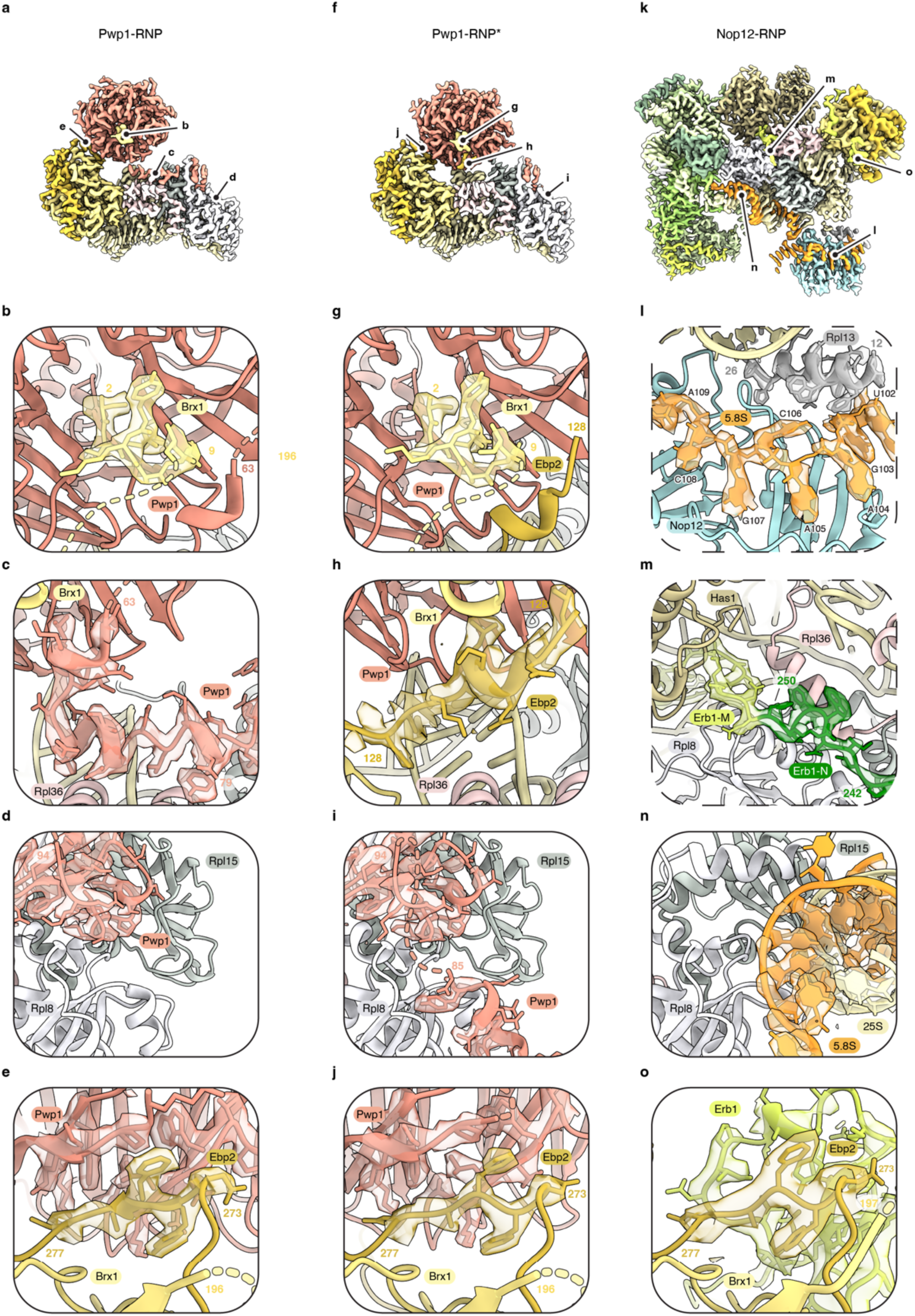
Representative cryo-EM densities and models of Pwp1 and Nop12 RNPs. Representative protein and RNA models with their corresponding cryo-EM density maps. (a, f, k) Composite cryo-EM maps of the Pwp1 RNP (a), Pwp1 RNP* (f), and Nop12-RNP (k) intermediates colored by protein chains. Labels indicate the positions of the remaining panels. (b, g) Close-up views of the Brx1 N-terminal docked onto the Pwp1 β-propeller. (c, h) Close-up views of the Pwp1 internal loop and Ebp2 binding sites, illustrating the mutual exclusivity between these factors during the transition from Pwp1 RNP to Pwp1 RNP*. (d, i, n) Detailed views of the Pwp1-Rpl8 interface. Pwp1 prevents the premature docking of the Rpl8 N-terminal and sterically obstructs the 25S-5.8S rRNA duplex (H10) as observed in Nop12-RNP. (e, j, o) Zoomed-in view of the Ebp2 binding interface. Ebp2 interaction with Pwp1 in early intermediates is subsequently replaced by Erb1 in the Nop12-RNP. (l) Close-up view of the 5.8S rRNA binding site on Nop12 highlighting the sequence of identified RNA bases and the resides of Rpl13 stabilizing Nop12 docking to the particle. (m) Detailed view of Erb1 interactions with components of Nop12-RNP. The Erb1(1-250) N-terminal segment is colored in darker green.

**Extended Data Fig. 4.**
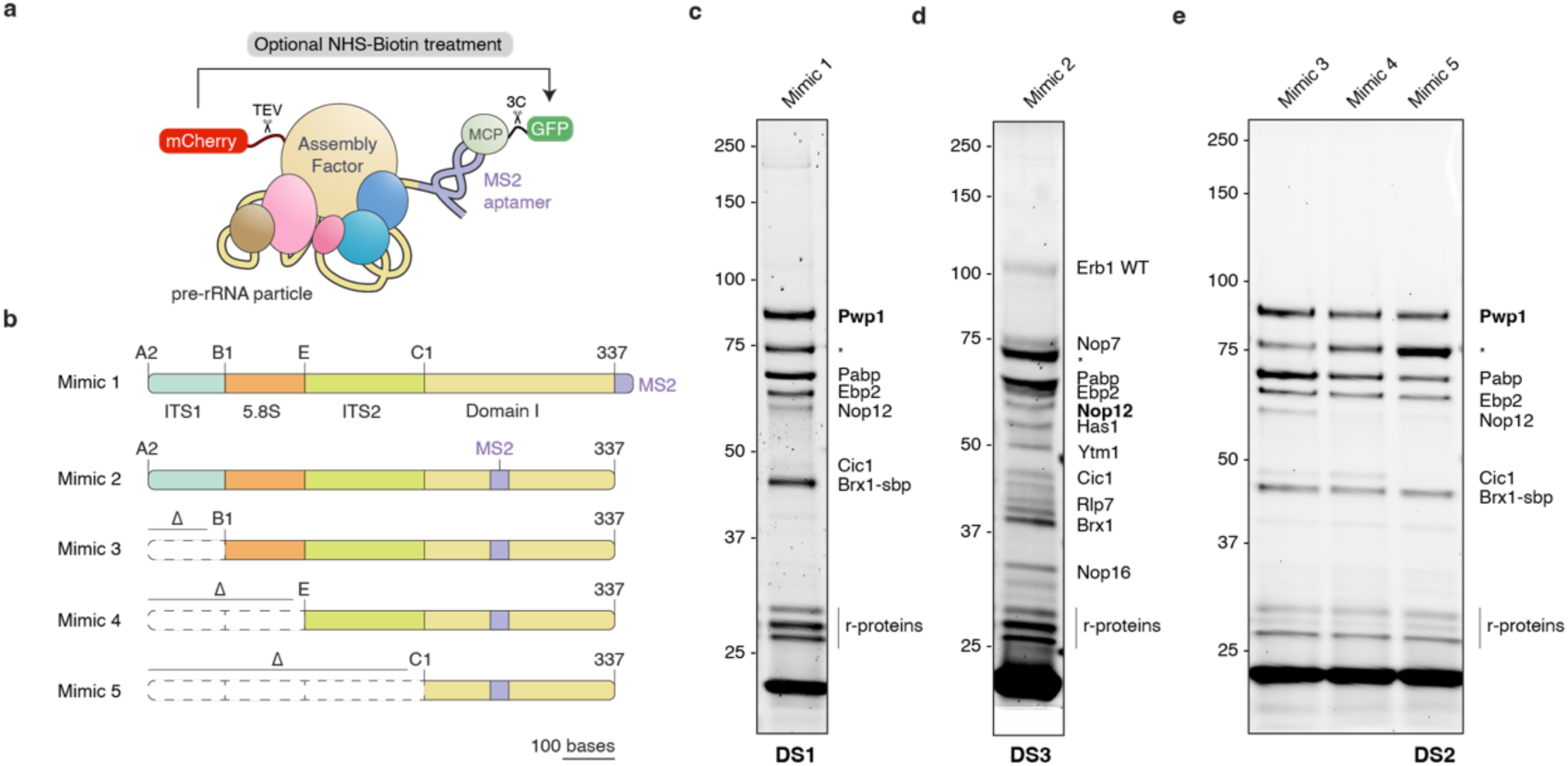
Purification and biochemical characterization of LSU assembly intermediates. (a) Schematic of the tandem affinity purification strategy for the isolation of pre-ribosomal intermediates. Sequential enrichment is facilitated by an mCherry-tagged assembly factor and an MCP-GFP fusion protein that specifically binds the MS2-tagged rRNA mimic. (b) rRNA mimics used in the study. Dashed lines indicate deleted segments. (c-e) SDS-PAGE analysis of purified assembly intermediates. Baits used for the primary pulldown are bolded. Combinations used for high-resolution structure determination are indicated (Datasets DS1-3).

**Extended Data Fig. 5.**
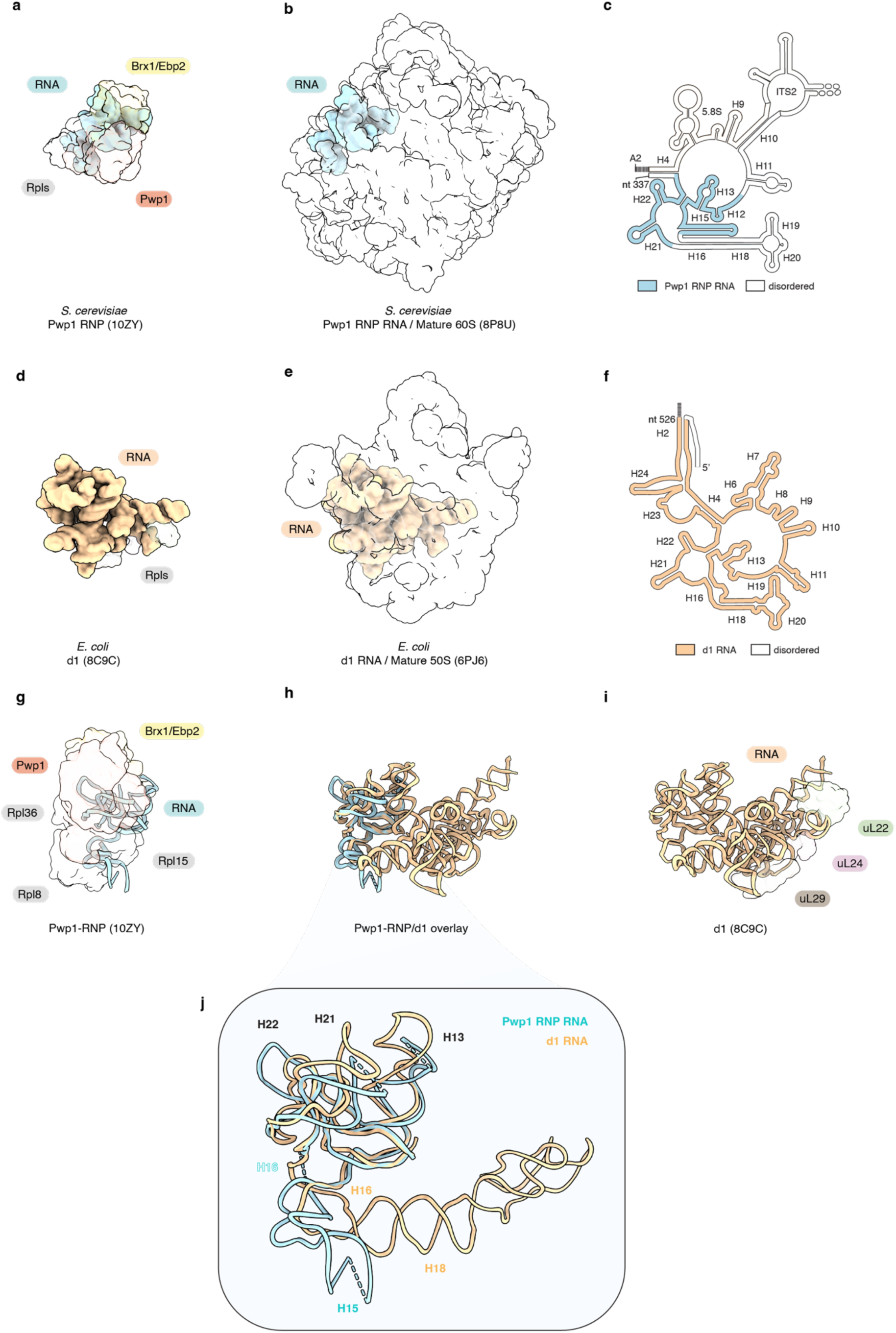
Comparison of the earliest experimentally determined LSU assembly intermediates for *S. cerevisiae* and *E. coli*. (a) Surface representation of *S. cerevisiae* Pwp1 RNP (PDB: 10ZY, this study) with proteins and rRNA visualized as transparent and solid surfaces respectively. (b) Overlay of Pwp1 RNP rRNA represented as blue surface with mature yeast 60S (PDB: 8P8U) represented as transparent surface. (c) Secondary structure of Pwp1 RNP rRNA with ordered RNA indicated in blue. (d-f) Corresponding analysis for the *E. coli* 50S assembly intermediate d1 (PDB: 8C9C). The d1 rRNA (gold) is compared to the mature 50S core (PDB: 6PJ6). (g-j) Comparative analysis of the early LSU assembly core in the yeast Pwp1 RNP (g) and the *E. coli* d1 intermediate (i). The structural superposition of both models (h) highlights significant differences in the size of ordered rRNA between the two intermediates. The inset (j) provides a high-resolution superposition of the conserved Domain I core, demonstrating the structural homology of helices H12, H13, H21, and H22. In contrast to the bacterial system, the eukaryotic H16-H20 region is incorporated during subsequent post-transcriptional assembly stages.

**Extended Data Fig. 6.**
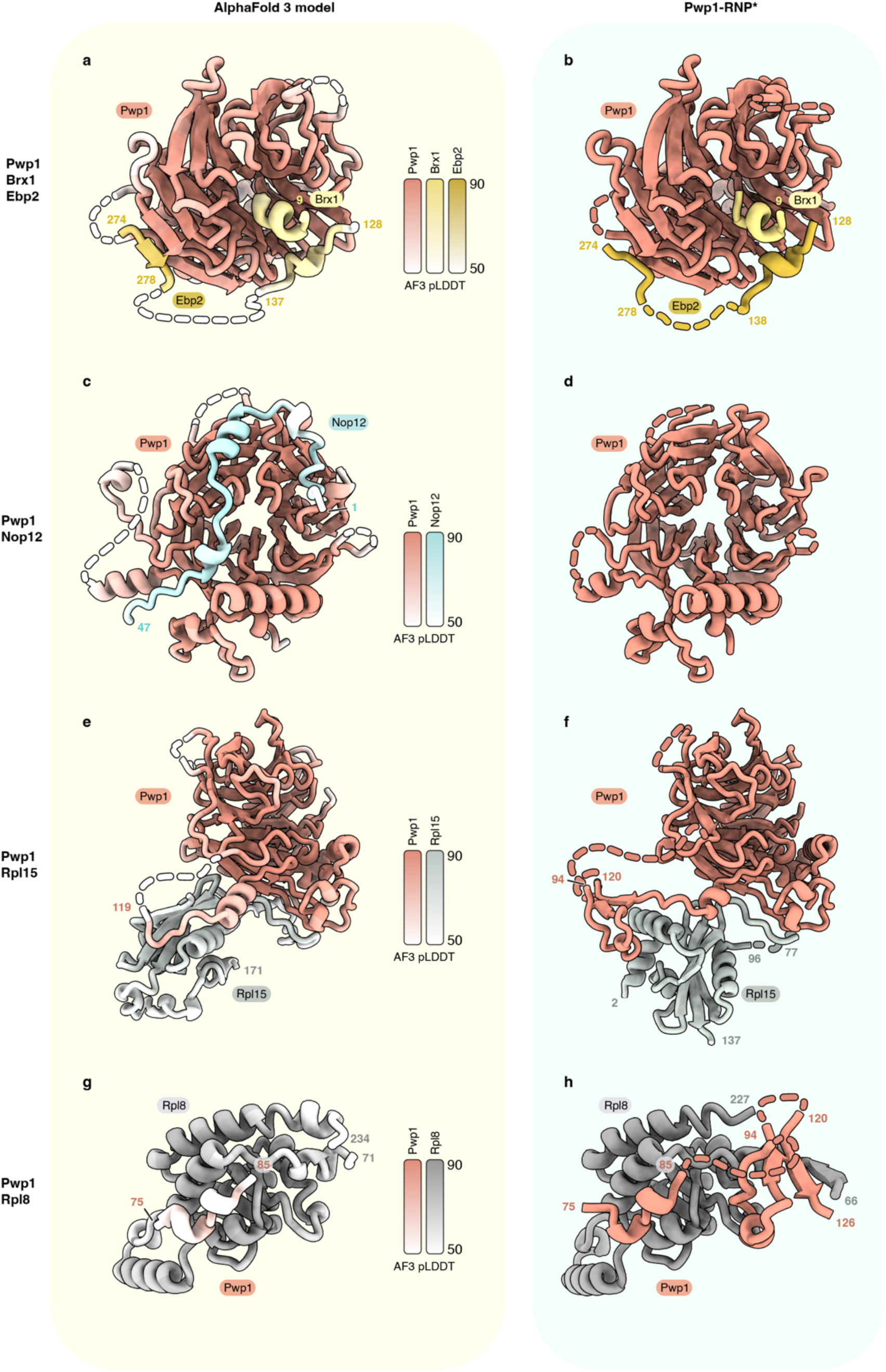
Comparison of selected AlphaFold3 predictions with experimentally determined models. (a, c, e, g) AlphaFold3 predicted models of Pwp1 in complex with Brx1/Ebp2 (a), Nop12 (c), Rpl15 (e), and Rpl8 (g). Models are colored by confidence score (pLDDT) as indicated by the scale bars. Full-length sequences were used for all predictions. (b, d, f, h) Respective experimentally determined protein-protein interactions in Pwp1 RNP*.

**Extended Data Fig. 7.**
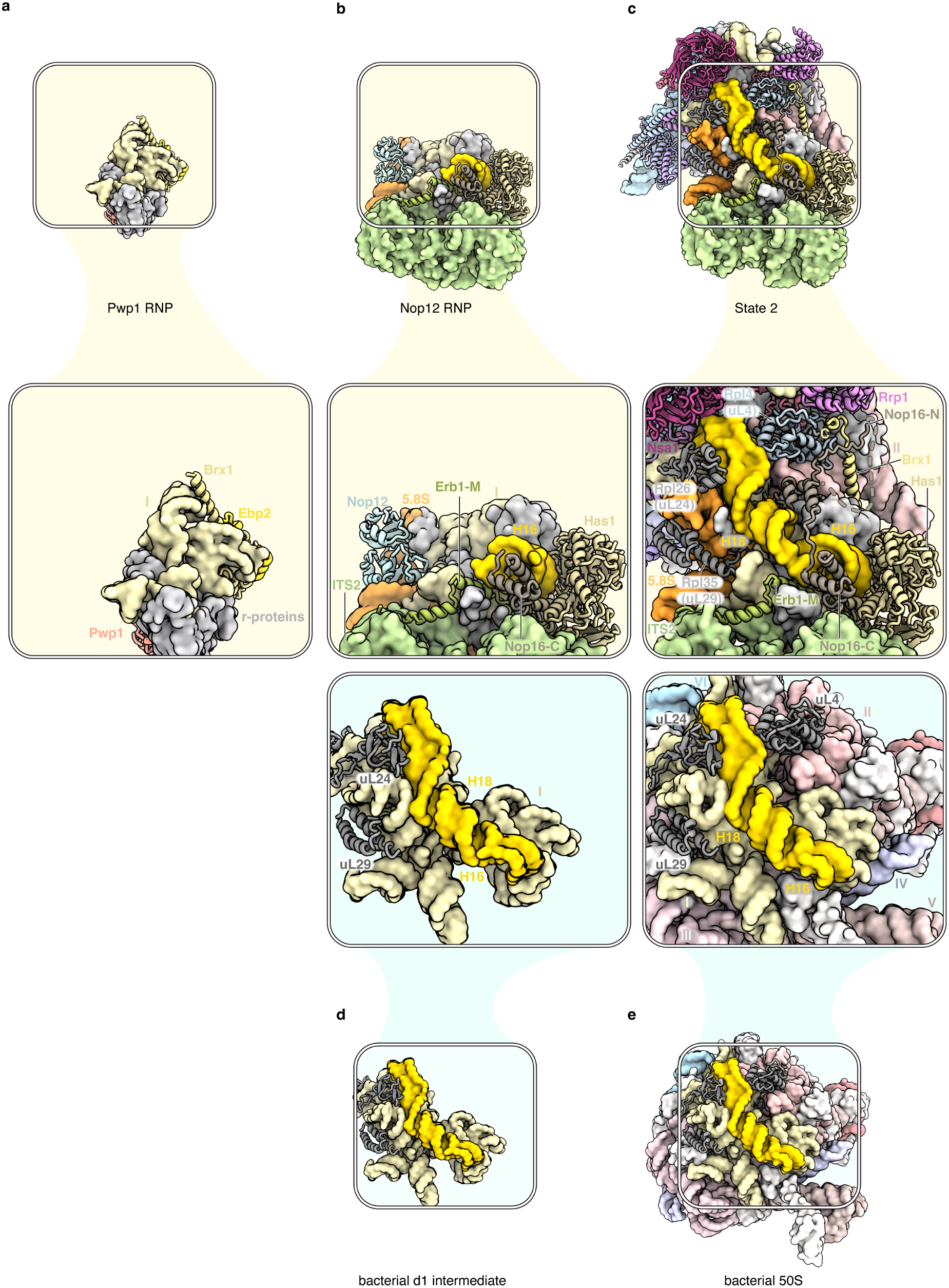
Stepwise maturation of the rRNA Helix 16 (H16) reveals divergent assembly pathways in eukaryotes and prokaryotes. (a-c), Sequential folding states of H16 across eukaryotic assembly intermediates. (a) The H16 region lacks discernible density in the earliest captured co-transcriptional intermediate, the Pwp1-RNP. (b) Initial stabilization of H16 (yellow) occurs within the Nop12-RNP through the concerted recruitment of Erb1-M, Has1, and Nop16. (c), Transition to post-transcriptional assembly stages, as visualized in the State 2 intermediate, allows H16-H20 to reach a near-mature conformation that stabilizes the Domain I-II interface. The recruitment of Nsa1, Rrp1, and ribosomal proteins uL4 and uL24 further clamps the region into its final orientation within the LSU core. (d, e), Comparative folding pathway of H16 in prokaryotes. (d) Structure of the bacterial d1 intermediate highlighting the H16-H20 region (yellow) in a near-mature conformation during the earliest stages of LSU assembly in prokaryotes. (e) The mature bacterial 50S subunit illustrates the final conserved position of the H16-H20 region within the ribosomal core. Comparison with eukaryotic states reveals that while the final H16-20 fold is conserved, the temporal order of its stabilization is delayed in eukaryotes until completion of the co-transcriptional assembly.

**Extended Data Table 1.**
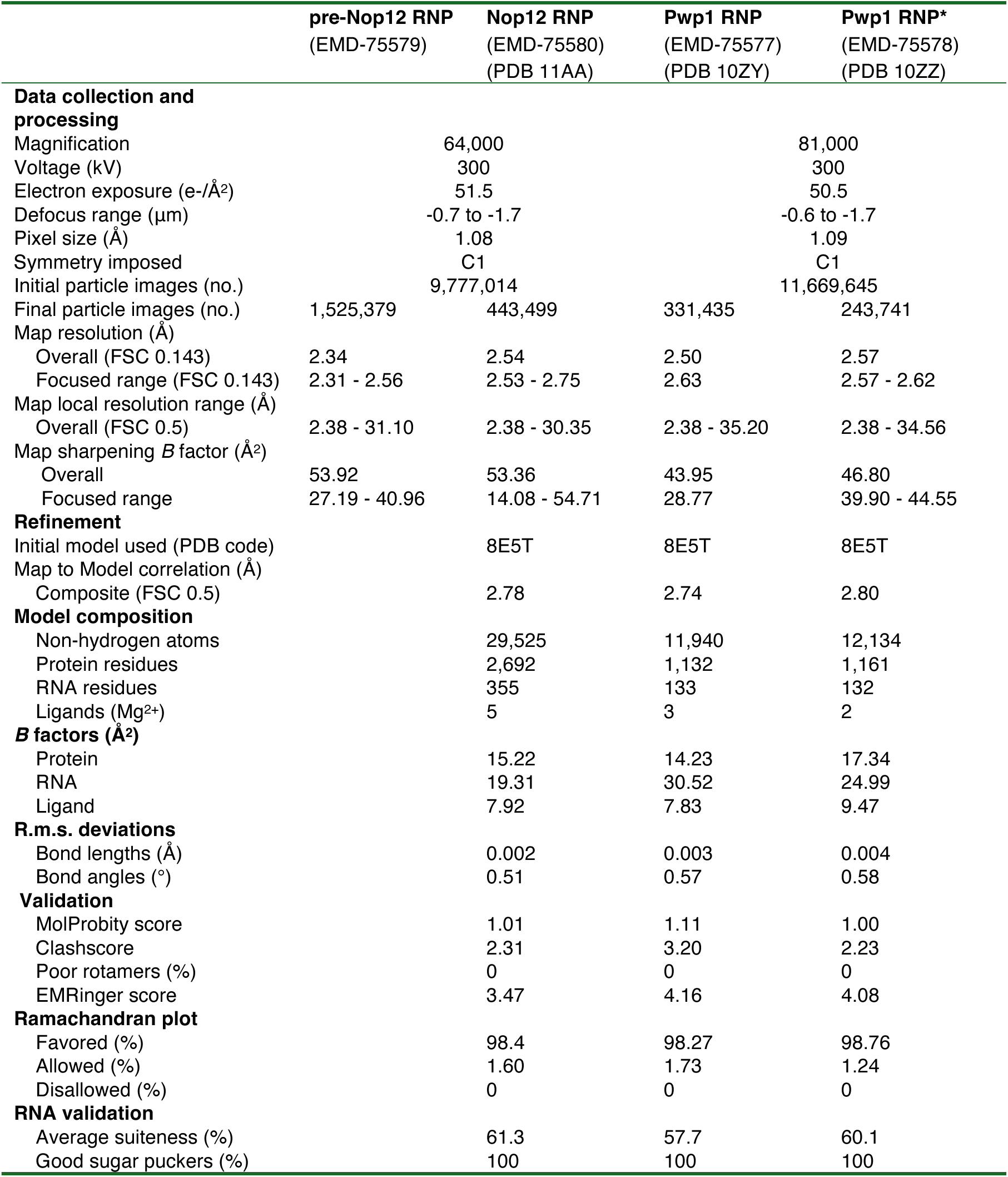
Cryo-EM data collection, refinement and validation statistics.

