## Supplementary Information for "Mechanism for the initiation of co-transcriptional pre-60*S* assembly"

### TABLE OF CONTENTS

Supplementary Fig. 1 Cryo-EM data processing of the Pwp1 A2→337 dataset. ____ 3

Supplementary Fig. 2 Cryo-EM data processing of the Pwp1 C1→337 dataset. ____ 5

Supplementary Fig. 3 Cryo-EM data processing of the Nop12 A2→337 dataset. ____ 7

Supplementary Fig. 4 Cryo-EM focused maps and composite reconstruction of pre-Nop12 RNP. ____ 9

Supplementary Fig. 5 Cryo-EM focused maps and composite reconstruction of Nop12 RNP. ____ 12

Supplementary Fig. 6 Cryo-EM data processing of merged Brx1 datasets. ____ 14

Supplementary Fig. 7 Cryo-EM focused maps and composite reconstruction of Pwp1 RNP and Pwp1 RNP*. ____ 16

Supplementary Fig. 8 Uncropped gel images. ____ 18

### Dataset 1: Pwp1-mCherry/MCP-GFP

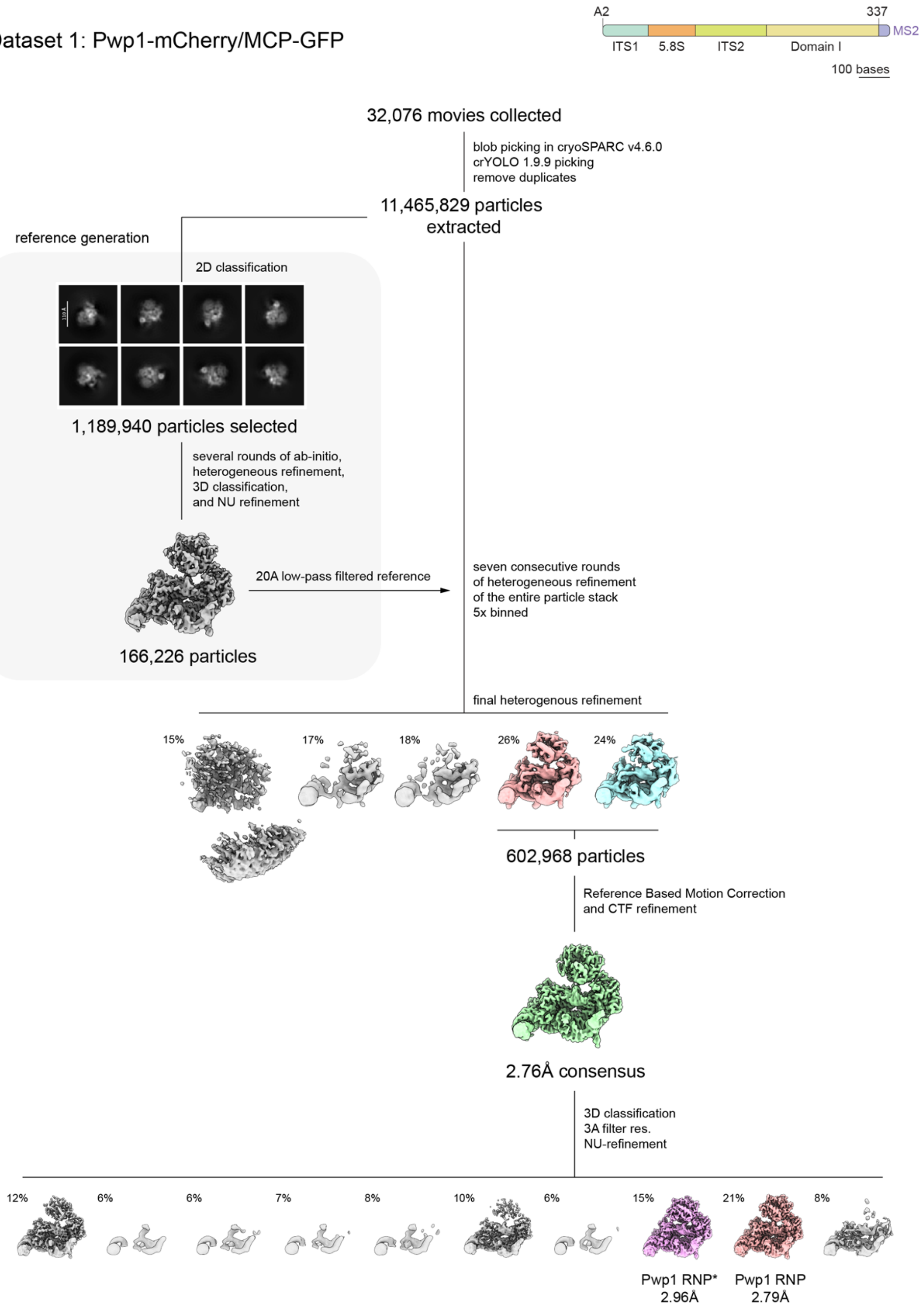

**Supplementary Fig. 1 Cryo-EM data processing of the Pwp1 A2→337 dataset (DS1).**

Schematic representation of the computational pipeline leading to the high-resolution reconstructions of the Pwp1-RNP and Pwp1-RNP* intermediates. Representative 2D class averages are shown. Samples were purified using mCherry-tagged Pwp1 and A2→337 rRNA mimic. Gold-standard Fourier shell correlation resolution values (GSFSC = 0.143) determined within cryoSPARC, are indicated below the respective density maps.

### Dataset 2: Pwp1-mCherry/MCP-GFP

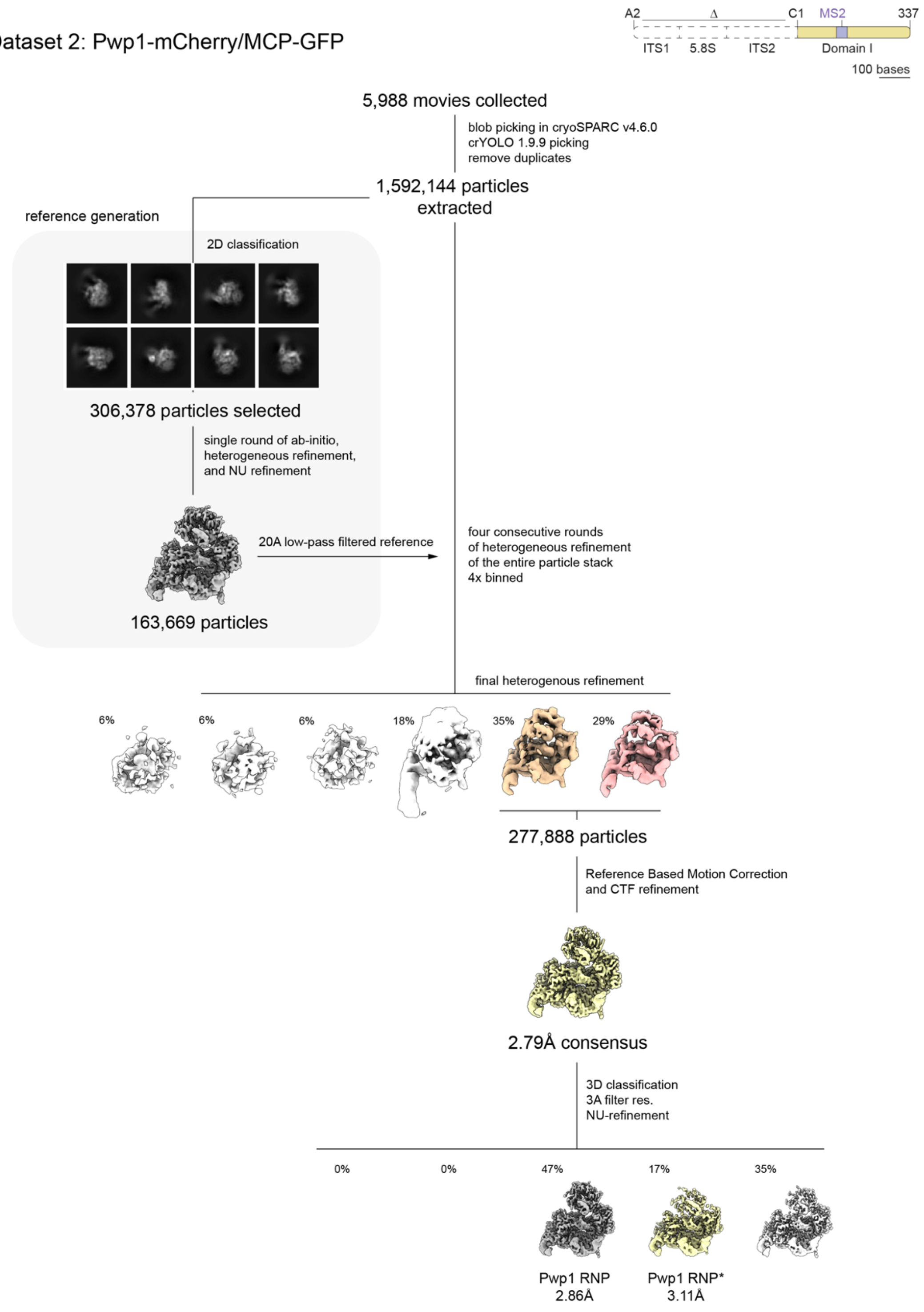

**Supplementary Fig. 2 Cryo-EM data processing of the Pwp1 C1→337 dataset (DS2).**

Schematic representation of the computational pipeline leading to the high-resolution reconstructions of the Pwp1-RNP and Pwp1-RNP* intermediates. Representative 2D class averages are shown. Samples were purified using mCherry-tagged Pwp1 and C1→337 rRNA mimic. Gold-standard Fourier shell correlation resolution values (GSFSC = 0.143) determined within cryoSPARC, are indicated below the respective density maps.

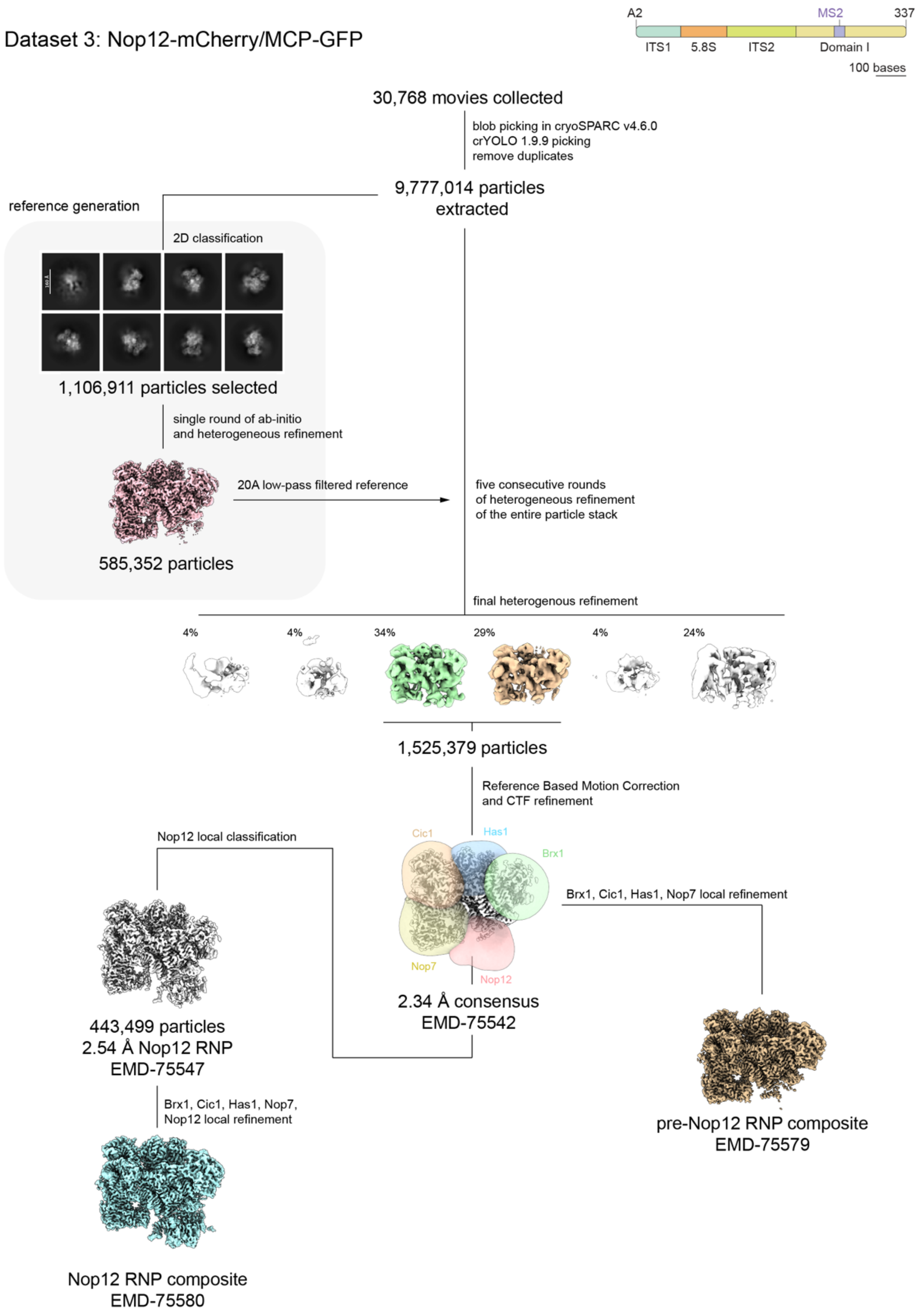
Dataset 3: Nop12-mCherry/MCP-GFP
A2
MS2
337
ITS1
5.8S
ITS2
Domain I
100 bases
30,768 movies collected
blob picking in cryoSPARC v4.6.0
crYOLO 1.9.9 picking
remove duplicates
9,777,014 particles
extracted
reference generation
2D classification
1,106,911 particles selected
single round of ab-initio
and heterogeneous refinement
20A low-pass filtered reference
585,352 particles
five consecutive rounds
of heterogeneous refinement
of the entire particle stack
final heterogenous refinement
4%
4%
34%
29%
4%
24%
1,525,379 particles
Reference Based Motion Correction
and CTF refinement
Nop12 local classification
Cic1
Has1
Brx1
Nop7
Nop12
Brx1, Cic1, Has1, Nop7 local refinement
443,499 particles
2.54 Å Nop12 RNP
EMD-75547
2.34 Å consensus
EMD-75542
Brx1, Cic1, Has1, Nop7,
Nop12 local refinement
pre-Nop12 RNP composite
EMD-75579
Nop12 RNP composite
EMD-75580

**Supplementary Fig. 3 Cryo-EM data processing of the Nop12 A2→337 dataset (DS3).**

Schematic representation of the computational pipeline leading to the high-resolution reconstructions of the pre-Nop12-RNP and Nop12-RNP intermediates. Representative 2D class averages are shown. Samples were purified using mCherry-tagged Nop12 and A2→337 rRNA mimic. Gold-standard Fourier shell correlation resolution values (GSFSC = 0.143) determined within cryoSPARC, are indicated below the respective density maps. The masks utilized for focused 3D classification and local refinement are shown overlaid on the consensus maps. The final composite maps are displayed.

a

pre-Nop12 RNP consensus EMD-75542

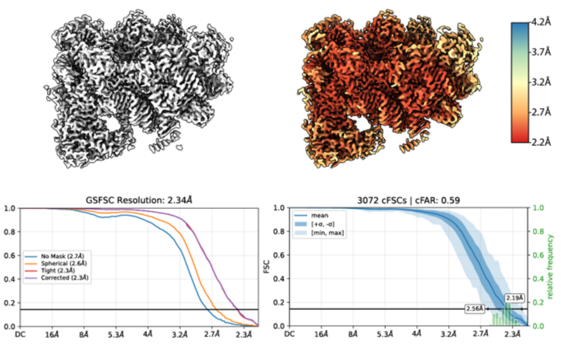


b

pre-Nop12 RNP composite EMD-75579

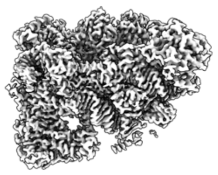

c

pre-Nop12 RNP local refinement Cic1 EMD-75543

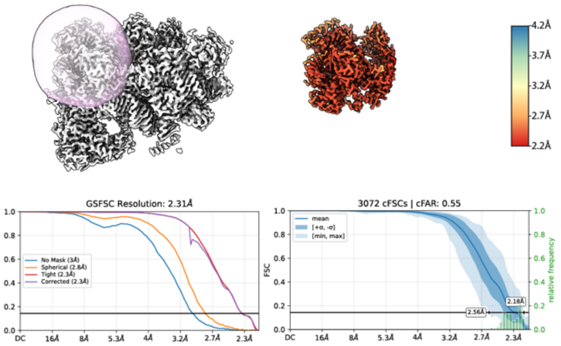


d

pre-Nop12 RNP local refinement Has1 EMD-75545

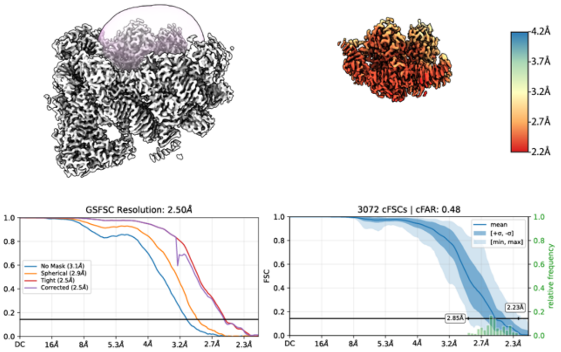


e

pre-Nop12 RNP local refinement Nop7 EMD-75546

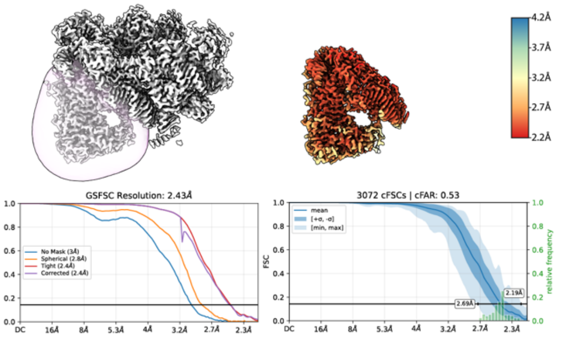


f

pre-Nop12 RNP local refinement Brx1 EMD-75544

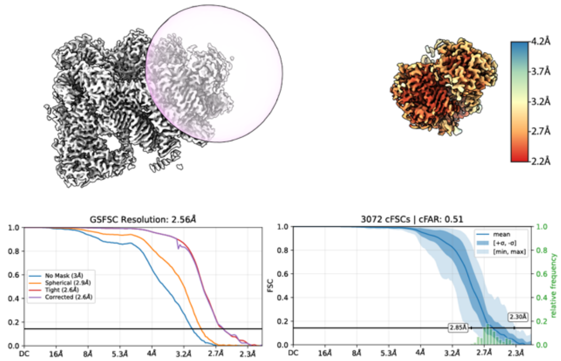

**Supplementary Fig. 4 Cryo-EM focused maps and composite reconstruction of pre-Nop12 RNP.**

(a) Consensus cryo-EM density (white) and corresponding local resolution map for the pre-Nop12-RNP. Gold-standard Fourier shell correlation (GSFSC) and conical FSC (cFSC) plots are provided below. (b) Final pre-Nop12 RNP composite map generated by combining the locally refined volumes. (c–f) Local refinements of Cic1 (c), Has1 (d), Nop7 (e), and Brx1 (f) contact sites. In each panel, the masks used for focused processing are highlighted (pink) on the consensus density. Corresponding local resolution maps, GSFSC curves and cFSC plots are provided. Resolution ranges are color-coded according to the scale bars.

**a**

Nop12 RNP consensus EMD-75547

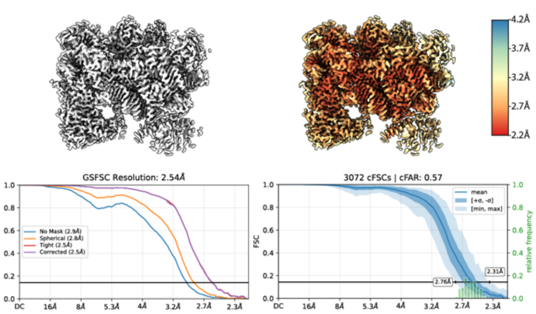


**b**

Nop12 RNP composite EMD-75580 PDB 11AA

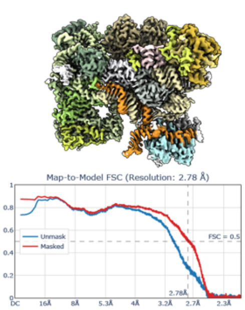


**c**

Nop12 RNP local refinement Cic1 EMD-75548

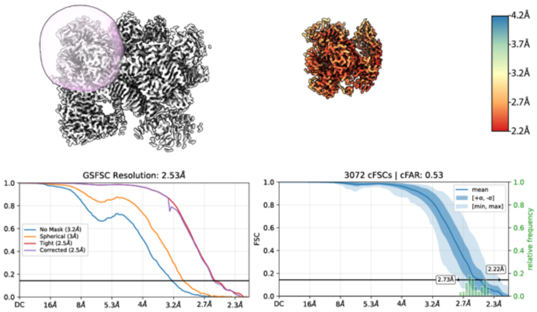


**d**

Nop12 RNP local refinement Has1 EMD-75550

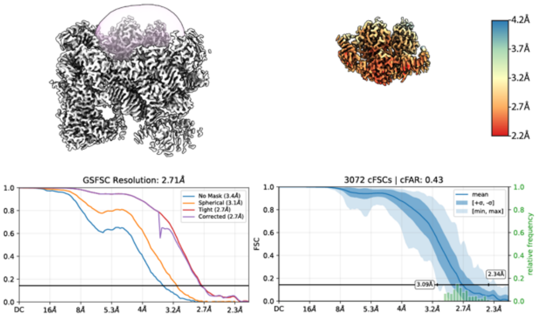


**e**

Nop12 RNP local refinement Nop7 EMD-75552

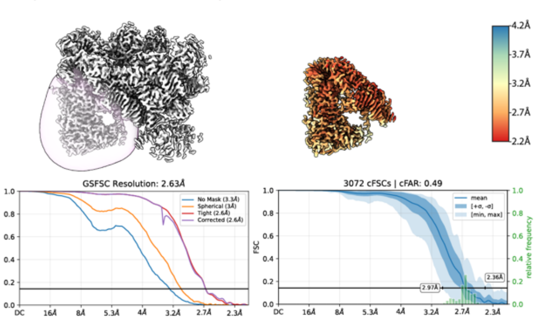


**f**

Nop12 RNP local refinement Brx1 EMD-75549

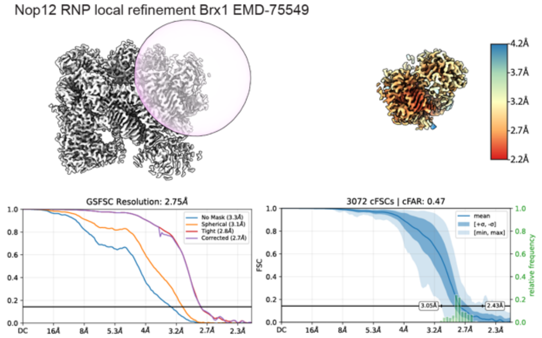


**g**

Nop12 RNP local refinement Nop12 EMD-75553

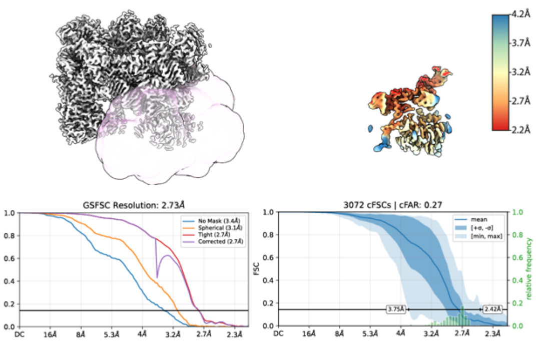

**Supplementary Fig. 5 Cryo-EM focused maps and composite reconstruction of Nop12 RNP.**

(a) Consensus cryo-EM density (white) and corresponding local resolution map for the pre-Nop12-RNP. Gold-standard Fourier shell correlation (GSFSC) and conical FSC (cFSC) plots are provided below. (b) Final Nop12 RNP composite map generated by combining the locally refined volumes, colored accordingly to the Nop12 RNP model. Map-to-model FSC is provided below. (c–g) Local refinements of Cic1 (c), Has1 (d), Nop7 (e), Brx1 (f), and Nop12 (g) contact sites. In each panel, the masks used for focused processing are highlighted (pink) on the consensus density. Corresponding local resolution maps, GSFSC curves and cFSC plots are provided. Resolution ranges are color-coded according to the scale bars.

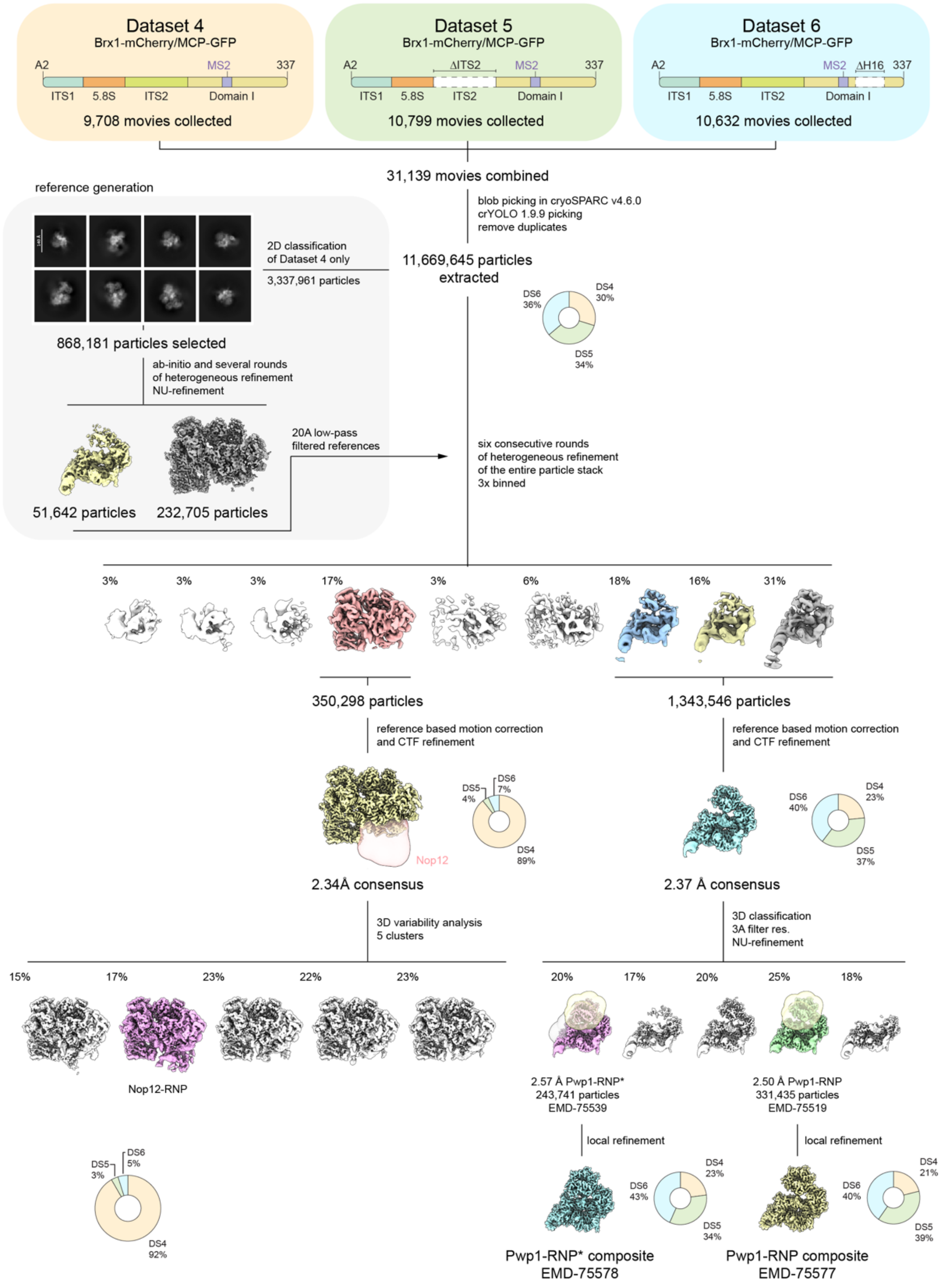
Dataset 4
Brx1-mCherry/MCP-GFP
A2
MS2
337
ITS1
5.8S
ITS2
Domain I
9,708 movies collected
Dataset 5
Brx1-mCherry/MCP-GFP
ΔITS2
10,799 movies collected
Dataset 6
Brx1-mCherry/MCP-GFP
ΔH16
10,632 movies collected
31,139 movies combined
blob picking in cryoSPARC v4.6.0
crYOLO 1.9.9 picking
remove duplicates
11,669,645 particles extracted
DS6 36%
DS4 30%
DS5 34%
reference generation
2D classification of Dataset 4 only
3,337,961 particles
868,181 particles selected
ab-initio and several rounds of heterogeneous refinement
NU-refinement
20A low-pass filtered references
51,642 particles
232,705 particles
six consecutive rounds of heterogeneous refinement of the entire particle stack
3x binned
3%
3%
3%
17%
3%
6%
18%
16%
31%
350,298 particles
1,343,546 particles
reference based motion correction and CTF refinement
reference based motion correction and CTF refinement
Nop12
DS5 4%
DS6 7%
DS4 89%
2.34Å consensus
DS6 40%
DS4 23%
DS5 37%
2.37 Å consensus
3D variability analysis
5 clusters
3D classification
3A filter res.
NU-refinement
15%
17%
23%
22%
23%
Nop12-RNP
DS5 3%
DS6 5%
DS4 92%
20%
17%
20%
25%
18%
2.57 Å Pwp1-RNP*
243,741 particles
EMD-75539
2.50 Å Pwp1-RNP
331,435 particles
EMD-75519
local refinement
local refinement
DS6 43%
DS4 23%
DS5 34%
DS6 40%
DS4 21%
DS5 39%
Pwp1-RNP* composite
EMD-75578
Pwp1-RNP composite
EMD-75577

**Supplementary Fig. 6 Cryo-EM data processing of merged Brx1 datasets (DS4-6).**

Schematic representation of the computational pipeline leading to the high-resolution reconstructions of the Pwp1-RNP and Nop12-RNP intermediates. Samples were purified using mCherry-tagged Brx1 in conjunction with either full-length (DS4), ΔITS2 (DS5), or ΔH16 (DS6) versions of A2→337 rRNA mimic. Representative 2D class averages are shown. Pie charts indicate the contribution of particles from each dataset to the final refined reconstructions. Gold-standard Fourier shell correlation resolution values (GSFSC = 0.143) determined within cryoSPARC, are indicated below the respective density maps. The masks utilized for focused 3D classification and local refinement are shown overlaid on the consensus maps. The final composite maps are displayed.

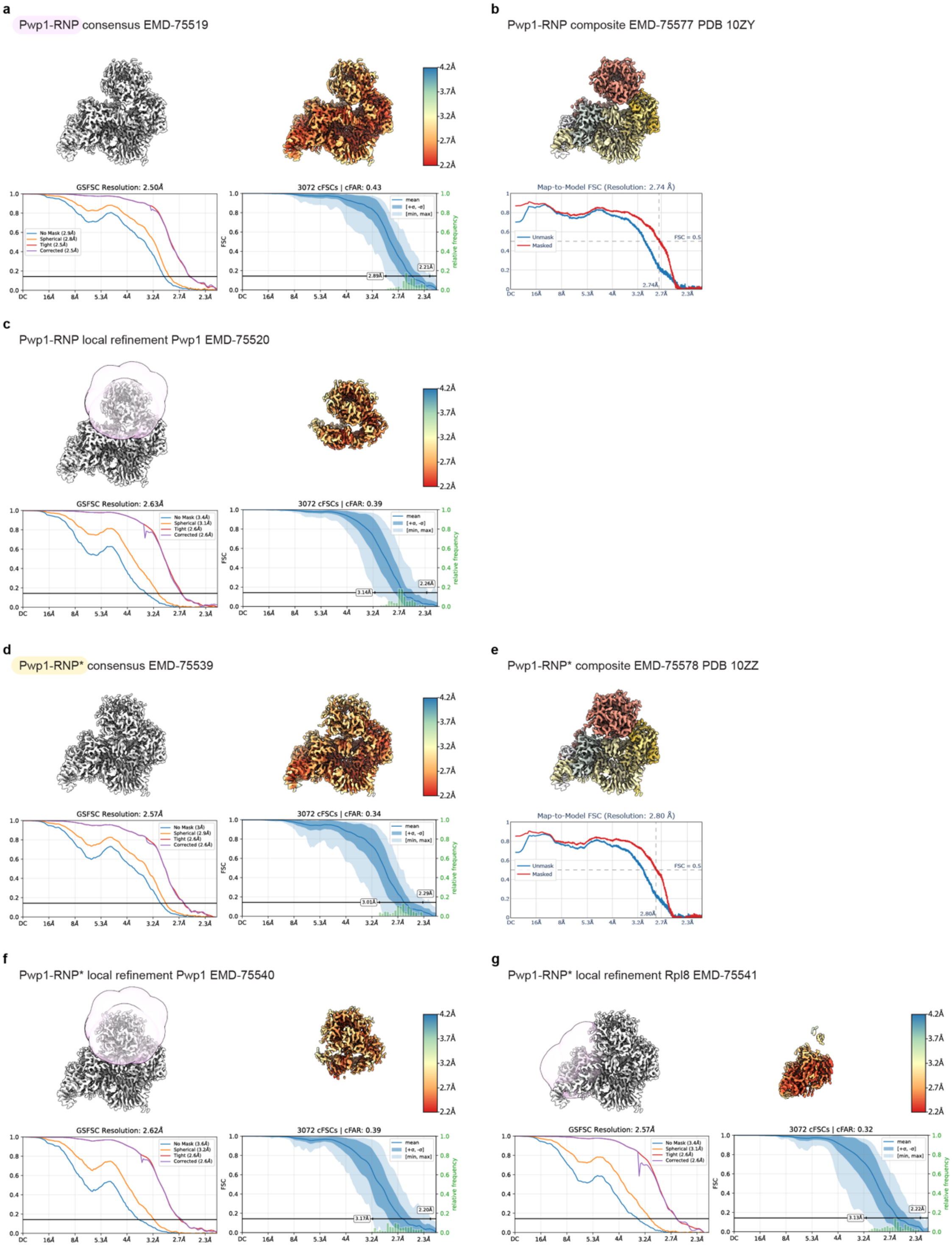
a
Pwp1-RNP consensus EMD-75519
GSFSC Resolution: 2.50Å
3072 cFSCs | cFAR: 0.43
b
Pwp1-RNP composite EMD-75577 PDB 10ZY
Map-to-Model FSC (Resolution: 2.74 Å)
c
Pwp1-RNP local refinement Pwp1 EMD-75520
GSFSC Resolution: 2.63Å
3072 cFSCs | cFAR: 0.39
d
Pwp1-RNP* consensus EMD-75539
GSFSC Resolution: 2.57Å
3072 cFSCs | cFAR: 0.34
e
Pwp1-RNP* composite EMD-75578 PDB 10ZZ
Map-to-Model FSC (Resolution: 2.80 Å)
f
Pwp1-RNP* local refinement Pwp1 EMD-75540
GSFSC Resolution: 2.62Å
3072 cFSCs | cFAR: 0.39
g
Pwp1-RNP* local refinement Rpl8 EMD-75541
GSFSC Resolution: 2.57Å
3072 cFSCs | cFAR: 0.32

**Supplementary Fig. 7 Cryo-EM focused maps and composite reconstruction of Pwp1 RNP and Pwp1 RNP*.**

(a, d) Consensus cryo-EM density (white) and corresponding local resolution map for the Pwp1-RNP (a) and Pwp1-RNP* (b). Gold-standard Fourier shell correlation (GSFSC) and conical FSC (cFSC) plots are provided below. (b, e) Final Pwp1 RNP (b) and Pwp1 RNP* (e) composite maps generated by combining the locally refined volumes, colored accordingly to models refined against the map. Map-to-model FSC is provided below. (c, f, g) Local refinements of Pwp1 site in Pwp1 RNP (c), and the Pwp1 (f) and Rpl8 (g) contact sites in Pwp1 RNP*. In each panel, the masks used for focused processing are highlighted (pink) on the consensus density. Corresponding local resolution maps, GSFSC curves and cFSC plots are provided. Resolution ranges are color-coded according to the scale bars.

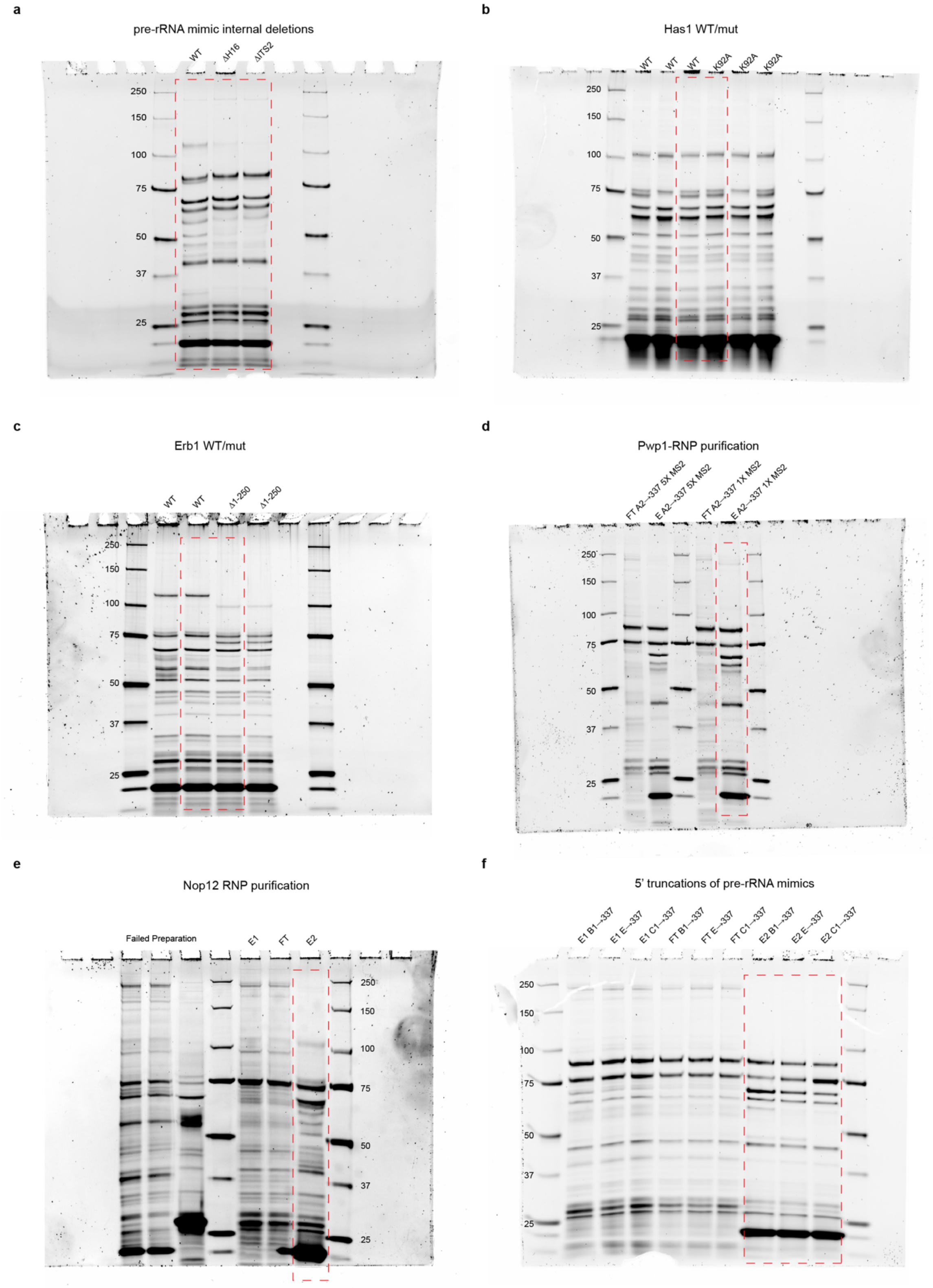
a
pre-rRNA mimic internal deletions
WT
ΔH16
ΔITS2
250
150
100
75
50
37
25
b
Has1 WT/mut
WT
WT
WT
K92A
K92A
K92A
250
150
100
75
50
37
25
c
Erb1 WT/mut
WT
WT
Δ1-250
Δ1-250
250
150
100
75
50
37
25
d
Pwp1-RNP purification
FT A2→337 5X MS2
E A2→337 5X MS2
FT A2→337 1X MS2
E A2→337 1X MS2
250
150
100
75
50
37
25
e
Nop12 RNP purification
Failed Preparation
E1
FT
E2
250
150
100
75
50
37
25
f
5' truncations of pre-rRNA mimics
E1 B1→337
E1 E→337
E1 C1→337
FT B1→337
FT E→337
FT C1→337
E2 B1→337
E2 E→337
E2 C1→337
250
150
100
75
50
37
25

**Supplementary Fig. 8 Uncropped gel images.**

Uncropped gel images relating to: (a) Fig. 3d, (b) Fig. 3e, (c) Fig. 3f, (d) Extended Data Fig. 3c (e) Extended Data Fig. 3d, (f) Extended Data Fig. 3e. Red dashed lines show approximate cropping dimensions used in related figures.
